# Crosstalk between Gut Sensory Ghrelin Signaling and Adipose Tissue Sympathetic Outflow Regulates Metabolic Homeostasis

**DOI:** 10.1101/2023.11.25.568689

**Authors:** M. Alex Thomas, Xin Cui, Liana R. Artinian, Qiang Cao, Jia Jing, Felipe C. Silva, Shirong Wang, Jeffrey M. Zigman, Yuxiang Sun, Hang Shi, Bingzhong Xue

**Affiliations:** Department of Biology, Georgia State University, Atlanta, GA; Department of Internal Medicine, University of Texas Southwestern Medical Center, Dallas, TX; Department of Nutrition, Texas A & M University, College Station, TX

## Abstract

The stomach-derived orexigenic hormone ghrelin is a key regulator of energy homeostasis and metabolism in humans. The ghrelin receptor, growth hormone secretagogue receptor 1a (GHSR), is widely expressed in the brain and gastrointestinal vagal sensory neurons, and neuronal GHSR knockout results in a profoundly beneficial metabolic profile and protects against diet-induced obesity (DIO) and insulin resistance. Here we show that in addition to the well characterized vagal GHSR, GHSR is robustly expressed in gastrointestinal sensory neurons emanating from spinal dorsal root ganglia. Remarkably, sensory neuron GHSR deletion attenuates DIO through increased energy expenditure and sympathetic outflow to adipose tissue independent of food intake. In addition, neuronal viral tract tracing reveals prominent crosstalk between gut non-vagal sensory afferents and adipose sympathetic outflow. Hence, these findings demonstrate a novel gut sensory ghrelin signaling pathway critical for maintaining energy homeostasis.

## Introduction

Prolonged energy intake that exceeds energy expenditure can result in obesity and markedly increases the risk for secondary health consequences such as type 2 diabetes, cardiovascular disease, and cancer (*1*). As such, there has been considerable effort to uncover the mechanisms regulating energy homeostasis. Previous research has primarily focused on energy intake regulation by the brain (*2*). Endocrine receptors that control energy homeostasis also are widely distributed in peripheral sensory neurons including gastric afferents that detect nutrients or distension (*3*), and these afferents are sufficient to regulate adipose tissue metabolism (*4*). As obesity incidence continues to increase worldwide, a comprehensive understanding of the peripheral mechanisms controlling ingestive behaviors and energy homeostasis is critically needed to provide more accessible pharmacological interventions independent of the brain and blood brain barrier.

The stomach-derived orexigenic hormone ghrelin (*5*), is a key mediator of metabolic homeostasis in humans due to its regulation of food intake, gut motility, energy expenditure, nutrient partitioning and glycemia (*6-9*). Circulating ghrelin levels rise pre-prandially (i.e. during fasting) and decrease post-prandially, and the rise and fall of ghrelin concentrations is suggested to contribute to normal energy homeostasis and overall metabolic health (*10, 11*). Interestingly, cold challenge also increases plasma ghrelin levels in humans (*12, 13*), and exogenous ghrelin decreases core body temperature and attenuates brown adipose tissue (BAT) thermogenesis (*14*), suggesting ghrelin signaling is also important during cold-induced adaptation responses. The ghrelin receptor, growth hormone secretagogue receptor 1a (GHSR), is widely expressed in the brain and on gastrointestinal vagal sensory neurons (*15-17*). In both humans and rodents, activation of GHSR markedly increases adiposity through increased food intake (*18, 19*), decreased energy expenditure, and reduced fatty acid utilization (*8, 20*). In contrast, GHSR deletion or blockade prevents diet-induced obesity (DIO), ghrelin-induced ingestive behaviors (*19, 21, 22*), and significantly increases energy expenditure and thermogenic capacity (*23-25*), and insulin sensitivity (*26*). However, GHSR restoration in the hypothalamic agouti-related neuropeptide (AGRP) neurons or the catecholaminergic neurons does not fully restore ghrelin’s effects suggesting additional central or peripheral ghrelin signaling is critical for metabolic control (*27, 28*).

*Ghsr* is expressed in the vagal afferent neurons (VAN) located in the nodose ganglia (*29*), and initial studies suggest that ghrelin-mediated regulation of energy expenditure, thermogenesis, gut motility, and glycemic control is dependent on intact gastric vagal afferents (*30-32*). However, vagotomized animals have normal food intake responses to an exogenous ghrelin challenge and normal adaptation to cold challenge (*33, 34*). In addition, a recent study using adeno-associated virus (AAV)-mediated *Ghsr* knockdown in the nodose ganglia in rats resulted in increased body weight over time (*29*), opposite to obesity resistance observed in whole body and neuronal *Ghsr* deficient mice (*21, 23*). On the other hand, non-vagal sensory neurons emanating from spinal dorsal root ganglia (DRG) innervate the gut at all levels (*35*), and total sensory neuron ablation following capsaicin treatment impairs adipose tissue metabolism, cold-induced thermogenic responses, and ghrelin’s gastric effects (*36-39*), suggesting that non-vagal DRG sensory neurons are important for metabolic control by ghrelin.

Here we show that DRG sensory neurons innervating gastrointestinal organs robustly express GHSR. Sensory neuron GHSR deletion attenuates DIO through increased energy expenditure and adipose tissue sympathetic outflow independent of food intake. Neuronal viral tract tracing further reveals prominent crosstalk between gut non-vagal sensory afferents and adipose sympathetic outflow. Hence, these findings demonstrate a novel gut sensory ghrelin signaling pathway critical for maintaining energy homeostasis.

## Results

### GHSR is expressed in vagal and DRG neurons that project to the gastrointestinal system

It was reported that cold challenge increases plasma ghrelin levels in humans (*12, 13*). Interestingly, we also found that cold exposure significantly increased plasma ghrelin levels in mice (**Figure 1A**). In addition, while both fasting and cold exposure significantly increased arcuate hypothalamus GHSR expression, only cold exposure significantly increased DRG GHSR expression (**Figures 1B-C**). Our data suggest that GHSR is expressed in DRG and that DRG GHSR may be potentially involved in regulating energy homeostasis under conditions of energetic challenge, such as cold exposure.

**Figure 1.**
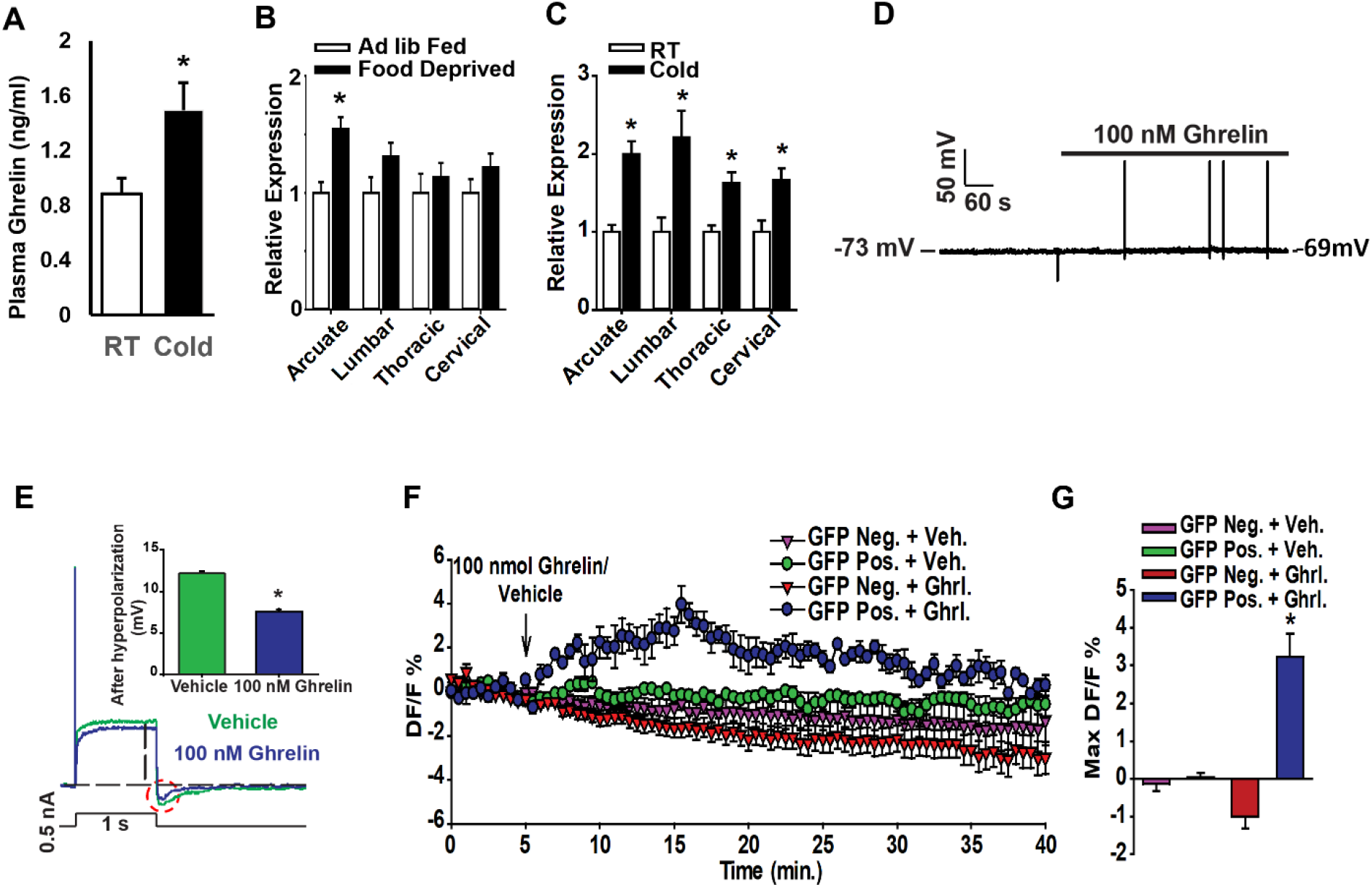
GHSR is expressed in DRG neurons. (A) Plasma acyl-ghrelin levels in 3-month-old male C57BL/6J mice after a 7-day cold exposure. n = 4 mice/group. (B) *Ghsr* expression in arcuate hypothalamus or lumbar, thoracic and cervical level DRGs in *ad lib*-fed and overnight food deprived mice; n = 6 mice/group. (C) *Ghsr* expression in arcuate hypothalamus or lumbar, thoracic and cervical level DRGs at room temperature or in response to 1-day 4°C cold exposure; n = 6 mice/group. (D) Representative continuous recording (current clamp mode) of resting membrane potential and spontaneous firing from GHSR-GFP-positive DRG neurons before and after ghrelin treatment. Ghrelin (100nM) depolarized membrane potential (+3.65 +/- 1.82 mV) (n=6 neurons) and stimulated spontaneous firing of action potentials (6/6 neurons tested). (E) Representative recording of voltage responses to an injection of 500pA current in the current clamp mode from GHSR-GFP-positive DRG neuron given either vehicle (green trace) or 100nM ghrelin (blue trace). Ghrelin suppressed the amplitude of the afterhyperpolarization (AHP) that is marked with dashed circle. (F) Average change in intracellular calcium in GFP-positive and GFP-negative DRG neurons given either 100 nM ghrelin (Ghrl.) or vehicle (Veh.); n = 5-12 mice/group. (G) Maximum change in intracellular calcium calculated from (D); n = 5-12 mice/group. Data presented as mean ± SEM. *, p<0.05 as measured by two-tailed unpaired t-test or one-way ANOVA with Bonferonni post-hoc analysis.

We then sought to identify gastrointestinal organs innervated by non-vagal DRG sensory neurons by injecting the retrograde neuronal tracer Fast Blue into different visceral organs including stomach, and small and large intestines. We found significant DRG innervation exists between T12 and T5 levels for the stomach, small intestine, and large intestine (**Figures S1A**). We subsequently investigated whether these DRG neurons that innervate the gastrointestinal organs express GHSR by injecting Fast Blue into the stomach, small and large intestines of GHSR-Tau-GFP reporter mice (*40*). We found extensive GFP and Fast Blue double labeled neurons in DRG ganglia that innervate the gastrointestinal organs (**Figures S1B**). Quantitatively, we observed GFP and Fast Blue colocalization in DRG neurons that project to the stomach (**Figure S1C**), small intestine (**Figure S1D**), and large intestine (**Figure S1E**). Within these neurons, 10%-40% GHSR-GFP neurons were co-labeled with Fast Blue and 20%-60% Fast Blue neurons were co-labeled with GHSR-GFP. These data suggest that out of the DRG GHSR-positive neurons, around 10-40% innervates stomach, small or large intestine; whereas out of DRG neurons innervate these gastrointestinal organs, between 20-60% are GHSR-positive.

We next confirmed that GHSR in these neurons was functional using cultured GHSR-tau-GFP DRG neurons *in vitro*. We found that 100nM ghrelin depolarized (+3.65 +/- 1.82 mV) and elicited spontaneous firing in GFP-positive DRG neurons (**Figure 1D**), similar to ghrelin’s effect on hypothalamic neuropeptide Y/agouti-related neuropeptide (NPY/AGRP) neurons (*20, 41, 42*). In addition, ghrelin suppressed afterhyperpolarization (AHP) after a single action potential evoked by a depolarizing current injection (500 pA, 1s) (**Figure 1E**), which has been shown to be mediated by calcium-activated potassium channels and contributes to DRG neuron excitability (*43*). Further, we also observed a marked increase in intracellular calcium in GFP-positive DRG neurons after 100nM ghrelin treatment (**Figures 1F-G**), similar to previous report (*44*). Importantly, this effect was absent in GFP-positive neurons receiving vehicle control and GFP-negative neurons receiving either 100nM ghrelin or vehicle control (**Figures 1F-G**).

### Sensory neuron GHSRs are activated by energetic challenges

We next tested whether these GHSR-positive DRG neurons could be activated by energetic challenges. We chose overnight food deprivation and acute (24 h) cold exposure challenges as these elicit a robust increase in circulating ghrelin (**Figure 1A**) (*10, 13*), and examined whether food deprivation or cold exposure activated DRG and vagal GHSR-positive neurons *in vivo* with GHSR-tau-GFP mouse line using triple labeling for GFP, Fast Blue, and the neuronal activity marker cFos (*45*) (**Figures 2A**). We found that whereas neither fasting nor cold exposure significantly affected the total number of GFP-positive DRG neurons (**Figures S1F-G**), both food deprivation and cold exposure significantly increased the activity of GHSR-positive DRG neurons projecting to stomach (**Figures 2B-C**) and small intestine (**Figures 2D-E**) as shown by increased number of neurons with GFP, Fast Blue and cFos triple labeling (**Figures 2B-E**). Similar results were observed when the data was expressed as percentage of neurons with GFP, Fast Blue and cFos triple labeling (**Figures S2A-D**), and this activation was present in both the nodose ganglia and T12-T5 DRG levels.

**Figure 2.**
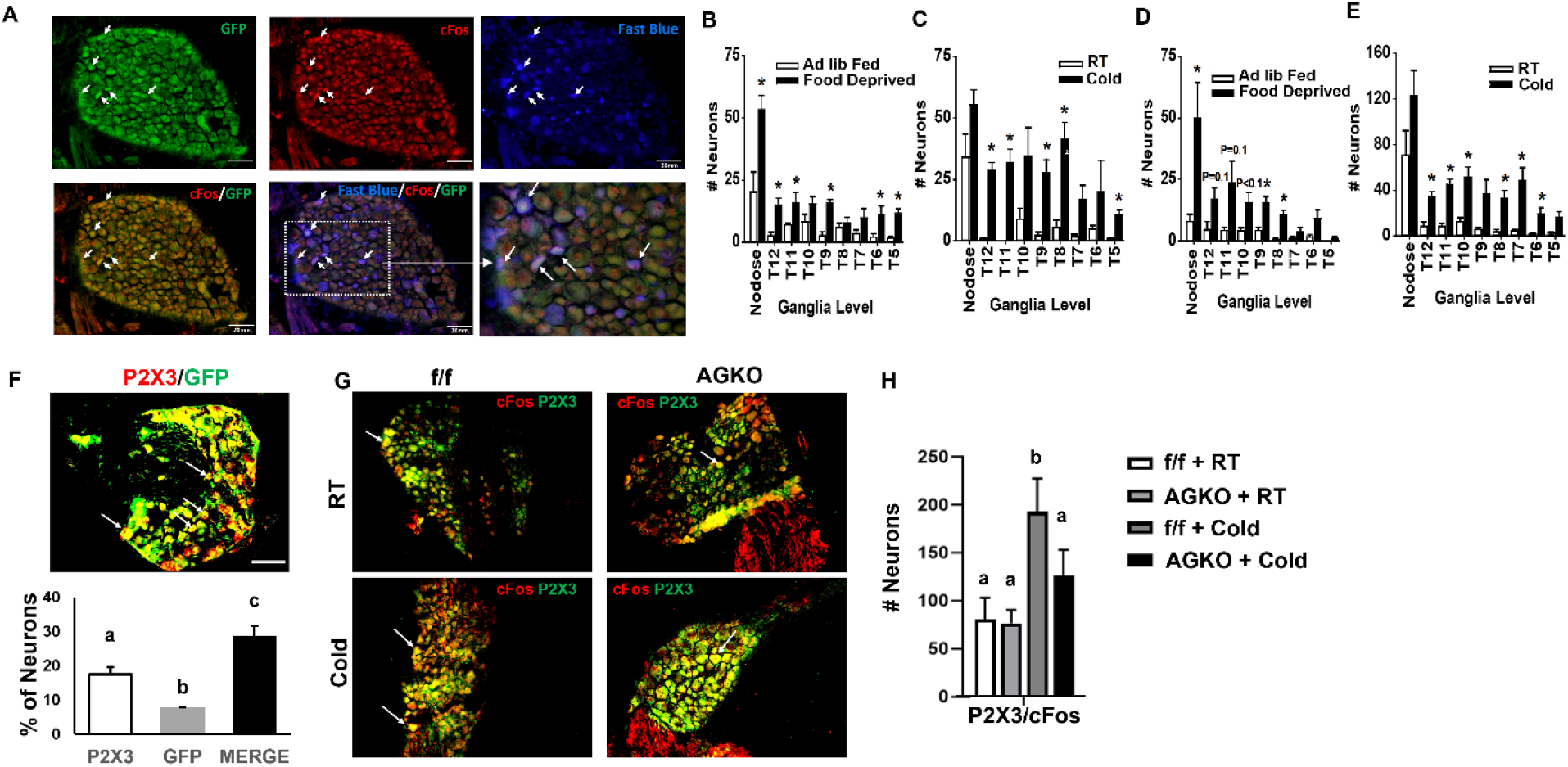
DRG GHSR-expressing neurons are regulated by energetic challenges. (A) Representative GFP, cFos, Fast Blue labeling, GFP and cFos double-labeling, and GFP/cfos/Fast Blue triple-labeling at T10 DRG level in stomach Fast Blue-injected mice following 1-day 4°C cold exposure. (B) Number of triple-labeled (GFP, cFos, and Fast Blue) neurons from nodose and T12-T5 level DRGs in stomach Fast Blue-injected mice following overnight food deprivation; n = 4 mice/group. (C) Number of triple-labeled (GFP, cFos, and Fast Blue) neurons from nodose and T12-T5 level DRGs in stomach Fast Blue-injected mice following 1-day 4°C cold exposure; n = 4 mice/group. (D) Number of triple-labeled (GFP, cFos, and Fast Blue) neurons from nodose and T12-T5 level DRGs in small intestine Fast Blue-injected mice following overnight food deprivation; n = 4 mice/group. (E) Number of triple-labeled (GFP, cFos, and Fast Blue) neurons from nodose and T12-T5 level DRGs in small intestine Fast Blue-injected mice following 1-day 4°C cold exposure; n = 4 mice/group. (F) Representative images and percentage of P2X3, GFP and P2X3/GFP double-labeled neurons at T10 DRG levels in GHSR-Tau-GFP mice. n = 3 mice/group. (G) Representative images of P2X3/cFos double-labeled neurons at T10 DRG levels in f/f and AGKO mice after 1-day cold exposure. (H) Percentage of P2X3/cFos double-labeled neurons at T10 DRG levels in f/f and AGKO mice after 1-day cold exposure. n = 4 mice/group. Data presented as mean ± SEM. In (F) and (H), bars with different lowercase letters are statistically different from each other. * p<0.05 as measured by two-tailed unpaired t-test or one-way or two-way ANOVA with Bonferonni post-hoc analysis.

We have further explored the neuronal identity of these GHSR-expressing neurons in DRG. We found that GHSR-GFP-positive DRG neurons were rarely (<3%) immunoreactive for neurofilament-200 (NF200) a marker of primary sensory neurons that give rise to myelinated axons (*46*) (**Figure S3A**). Unmyelinated C fibers, most of which are nociceptors, can be broadly subdivided into two classes, ’peptidergic’ and ’non-peptidergic’ (*47, 48*). The peptidergic group expresses neuropeptides such as calcitonin gene-related peptide (CGRP) and requires nerve growth factor (NGF)/Tropomyosin receptor kinase A (TRKA) signaling for survival. In contrast, the non-peptidergic group lacks peptides, requires glial cell-derived neurotrophic factor (GDNF)/c-RET signaling for survival and binds the plant lectin IB4. The non-peptidergic subclass express the ATP-gated ion channel purinergic receptor P2X ligand-gated ion channel 3 (P2X3); whereas tyrosine hydroxylase (TH) also marks a subclass of non-peptidergic C-fibers that respond to low-threshold stimuli (C-LTMRs) (*47, 48*). Interestingly, we found that <6% of the GHSR-GFP-positive neurons were immunoreactive for CGRP, TRKA or TH (**Figures S3B-D**); whereas most of the GHSR-positive DRG neurons were immunoreactive for P2X3 (**Figure 2F**). Thus, our data suggest that the majority of GHSR-expressing neurons in DRG belong to unmyelinated non-peptidergic C-fiber class that expresses P2X3.

We then further studied whether these GHSR/P2X3 positive DRG neurons could be regulated by energic challenges such as cold exposure. For this purpose, we have generated sensory neuron-specific GHSR knockout mice. To accomplish this, *Syn1-Cre;Ghsr^f/f^* mice (*23*) were first bred with wild type C57BL/6J mice to obtain *Ghsr^f/f^* mice, which were then bred with *Advillin-Cre* mice (*49*) to generate *Advillin-Cre+/-;Ghsr^f/f^* mice (henceforth referred to as AGKO). AGKO mice had normal GHSR expression in the brain, including the hypothalamus and the ventral tegmental area (VTA), but >90% decrease in DRG and nodose ganglia GHSR expression compared to that of f/f mice, indicating GHSR deletion is specific to peripheral sensory neurons (**Figure S4A-B**). We then subjected AGKO mice and their f/f littermates to an overnight cold challenge and measured P2X3-expressing neuron activities via P2X3/cFos double staining in DRG. As shown in **Figure 2G-H**, cold exposure significantly increased number of P2X3/cFos double-labeled neurons in DRG of f/f mice; however, this increase was blunted in AGKO mice (**Figure 2G-H**).

### Sensory neuron GHSR knockout mice are resistant to diet-induced obesity due to increased energy expenditure and adipose tissue thermogenesis

To examine the endogenous role of sensory neuron GHSR signaling in regulating energy homeostasis, we studied the metabolic phenotypes of AGKO mice and their f/f littermates fed either a regular chow or a high fat diet. We found that plasma ghrelin level was not different between f/f and AGKO mice on regular chow diet (**Figure S5A**). In addition, there was no difference in body weight between f/f and AGKO mice on regular chow diet up to 5-month of age (**Figure S5B**). Interestingly, chow-fed aging 8-month-old AGKO mice exhibited lower body weight than that of age-matched f/f mice (**Figures S5B**). More careful examination of the animals revealed that there was no difference in body length (**Figures S5B**) and in interscapular brown adipose tissue (iBAT), epididymal adipose tissue (eWAT), inguinal adipose tissue (iWAT), or liver mass in chow-fed 3-month-old AGKO mice compared to that of f/f littermates (**Figures S5C**). Similarly, there was also no difference in percentage and total lean and fat mass between chow-fed f/f and AGKO mice at 5 months of age (**Figure S5D**). Interestingly, consistent with their lower body weight (**Figure S5B**), aging 8-month-old chow-fed AGKO mice showed significantly reduced percentage and total fat mass compared to their f/f littermates without any differences in lean mass (**Figure S5E**).

Ghrelin and GHSR are important in regulating food intake (*18, 19, 21*) and glucose homeostasis (*21, 28, 50, 51*). We found exogenous ghrelin stimulated comparable 30 minutes and total 2 hours food intake in 12-week-old f/f and AGKO mice (**Figure S6A**). In addition, there was also no difference in food intake evoked by food deprivation (**Figure S6B**) or measured in a 24-hour period (**Figure S6C**) between AGKO and f/f control littermates on chow. Further, there was no difference in glucose and insulin tolerance tests (GTT and ITT, respectively) (**Figures S6D-E**), and blood glucose and insulin levels (**Figures S6F-G**) between AGKO and f/f mice on the chow diet, when their body weight was not different, indicating sensory GHSR is not necessary in regulating food intake or glucose homeostasis.

Interestingly, we found that chow-fed 12-week-old AGKO and f/f control mice had slightly elevated energy expenditure at several time points during both daytime and nighttime (**Figure S7A**), while there was no difference in locomotor activity between AGKO and f/f mice during either daytime or nighttime (**Figure S7B**). In addition, uncoupling protein 1 (UCP1) protein level was slightly increased in iBAT and showed a trend of increase in iWAT (**Figures S7C-D**), which corresponds to smaller adipocytes and more UCP1 IHC staining in iBAT and iWAT (**Figures S7E-F**). Thus, our data suggest that the leaner phenotype in aging 8-month-old chow-fed AGKO mice may be a result of increased energy expenditure, but may not be due to any differences in food intake, and sensory GHSR may be involved in regulating brown/beige adipocyte thermogenesis and energy expenditure.

It was reported that HFD feeding for 12 weeks reduced plasma ghrelin levels and *Ghsr* expression in nodose ganglia and hypothalamus (*52*). However, we found that there was no difference in *Ghsr* expression in hypothalamus or DRG of C57BL/6J mice fed HFD for 4 weeks (**Figure S8A**), indicating the reduced Ghsr expression in nodose ganglia and hypothalamus observed by Naznin et al (*52*) may be due to a chronic adaptation to the HFD. Thus, to further test the role of sensory GHSR in the development of diet-induced obesity, we studied metabolic phenotypes of AGKO mice challenged with a HFD. As shown in **Figure S8B**, HFD-fed AGKO mice had comparable plasma ghrelin levels when compared to that of f/f mice. Interestingly, when fed a HFD, AGKO mice gained significantly less body weight and had overall lower fat mass than their *f/f* littermate control mice, indicating that AGKO mice are resistant to the development of diet-induced obesity (DIO) (**Figures 3A-B**). Tissue analysis revealed a significant decrease in liver and iWAT mass (**Figure 3C**), corresponding to smaller adipocytes in iBAT, iWAT and eWAT, as well as reduced lipid accumulation in liver (**Figure 3D**). Critically, we found the resistance to DIO in AGKO mice was independent of changes in food intake or fat absorption. Ten-to-twelve weeks old HFD-fed AGKO and f/f mice had comparable food intake following an exogenous ghrelin challenge (**Figure S9A**), an overnight food deprivation challenge (**Figure S9B**), as well as 24- hour total intake (**Figure S9C**). In addition, there was no difference in 24-hour fecal lipid content between HFD-fed AGKO and f/f (**Figure S9D**), suggesting sensory GHSR deficiency did not affect fat absorption.

**Figure 3.**
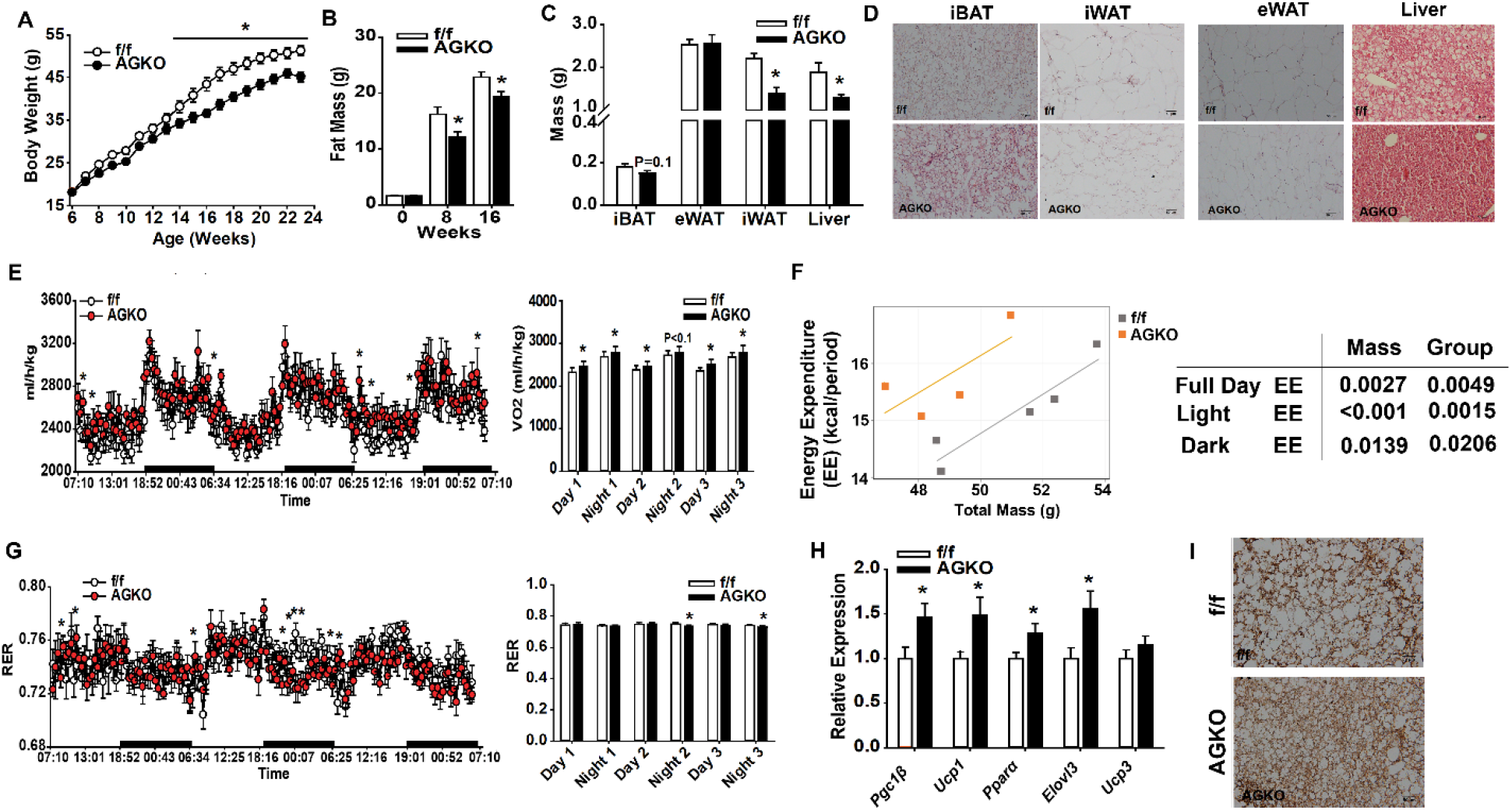
Sensory neuron GHSR knockout mice exhibit attenuated diet-induced obesity. (A) Weekly body mass measurements in f/f and AGKO mice fed 60% HFD; n = 15-17 mice/group. (B) Total fat mass measured by a Minispec LF90 TD-NMR analyzer in f/f and AGKO mice before the 60% HFD feeding (Week 0) and after 8 and 16 weeks of 60% HFD feeding; n = 15-17 mice/group. (C) Tissue weights following 17-week 60% HFD feeding in f/f and AGKO mice; n = 8 mice/group. (D) Representative H&E staining in iBAT, iWAT, eWAT and liver of f/f and AGKO mice fed HFD for 17 weeks. (E) Average oxygen consumption rate during daytime and nighttime in f/f and AGKO mice fed 60% HFD for 16 weeks; n = 7 mice/group. (F) ANCOVA analysis of energy expenditure data from f/f and AGKO mice fed with a HFD for 16 weeks. (G) Average respiratory exchange ratio (RER) during daytime and nighttime in f/f and AGKO mice fed 60% HFD for 16 weeks; n = 7 mice/group. (H) Relative expression of thermogenic genes in iBAT of f/f and AGKO mice fed 60% HFD for 17 weeks; n = 8 mice/group. (I) Representative images of iBAT UCP1 IHC staining in f/f and AGKO mice fed 60% HFD for 17 weeks. Data presented as mean ± SEM. *, p<0.05 versus f/f controls as measured by two-tailed unpaired t-test or two-way ANOVA with Bonferonni post-hoc analysis.

In contrast, HFD-fed AKO mice had significantly increased energy expenditure during both daytime and nighttime compared to that of f/f control mice (**Figure 3E**). Since AGKO mice were leaner than the f/f control mice, we further analyzed whether the differences in energy expenditure is dependent on their body weight difference by a regression-based analysis of covariance (ANCOVA) using the online software CalR (https://calrapp.org/) as previously described (*53*). By ANCOVA analysis, we found that the higher energy expenditure in AGKO mice in response to the HFD feeding is independent of the differences in body weight (**Figure 3F**). Importantly, this increase in energy expenditure was not due to changes in locomotor activity (**Figure S9E**). In addition, AGKO mice had significantly decreased respiratory exchange rate (RER) during nighttime hours indicating a preferential utilization of fatty acids as fuel (**Figure 3G**). Consistent with increased energy expenditure, we found a significant increase in thermogenic gene expression and UCP1 IHC staining in iBAT of HFD-fed AGKO mice compared to that of f/f control mice (**Figure 3H-I**). In addition, the expression of thermogenic genes, including the β3 adrenergic receptor (*β3ar*) and peroxisome proliferator activated receptor gamma (*Pparγ)*, was also significantly increased in both eWAT and iWAT (**Figure S9F-G**). Our data suggest that DIO resistance in AGKO mice is primarily due to increased energy expenditure, which is associated with upregulated thermogenic pathways in adipose tissue.

The significant reduction in body mass and adipocyte size suggests AGKO adipose depots are metabolically healthier than their f/f littermate controls. Indeed, we found significant decreases in the expression of inflammatory and fibrosis-related genes in iBAT and eWAT of HFD-fed AGKO mice (**Figures S10A-B**), indicating adipose tissue inflammation and fibrosis are significantly reduced in AGKO mice compared to that of f/f controls. Consistent with overall improved metabolic health, HFD-fed AGKO mice exhibited improved insulin sensitivity, as indicated by their lower fasting glucose and blood insulin levels (**Figures S10C-D**), lower glucose excursion response during GTT (**Figure S10E**) and an increased trend of glucose disposal during ITT (**Figure S10F**).

Ambient temperature poses a mild cold stress to the animals that results in sympathetic activation to maintain their body temperature, whereas under thermoneutrality (30°C) the sympathetic nervous system is minimally activated (*54-56*). To further investigate whether the leaner phenotype in AGKO mice is dependent on sympathetic activation, we also conducted HFD challenge in AGKO and f/f mice under thermoneutrality (30°C). Interestingly, we found that the DIO protective effects of AGKO disappeared in HFD-fed AGKO and f/f mice housed under thermoneutrality, as there was no difference in body weight between HFD-fed AGKO and f/f mice under thermoneutrality (**Figure S11A**). In addition, body composition analysis also showed that there was no difference in total fat mass and lean mass between AGKO and f/f mice when measured in mice prior to HFD challenge (6-week-old) (**Figure S11B**) and after 10 weeks on the HFD (**Figure S11C**). These data suggest that sensory GHSR regulation of energy homeostasis might be dependent on sympathetic activity.

### AGKO mice exhibit increased cold-induced thermogenesis

We next tested the thermoregulatory ability in AGKO and f/f mice during an acute (1-day) or chronic (7-day) cold challenge. We found that AGKO mice had significantly increased baseline body temperature when housed at ambient temperature (**Figures S12A-B**) and were able to normalize their body temperature more rapidly compared to that of f/f control mice in response to an acute cold challenge (**Figure 4A**). In addition, AGKO mice also had significantly higher core body temperature than that of f/f mice during a chronic 7-day cold challenge (**Figure 4B**). During the cold challenge, AGKO mice had significantly reduced body mass (**Figure S12C**), but this change was not explained by changes in food intake, as there was no difference in either acute food intake (**Figure S12D**) or total food intake across three days during the chronic cold exposure between AGKO and f/f mice (**Figure S12E**).

**Figure 4.**
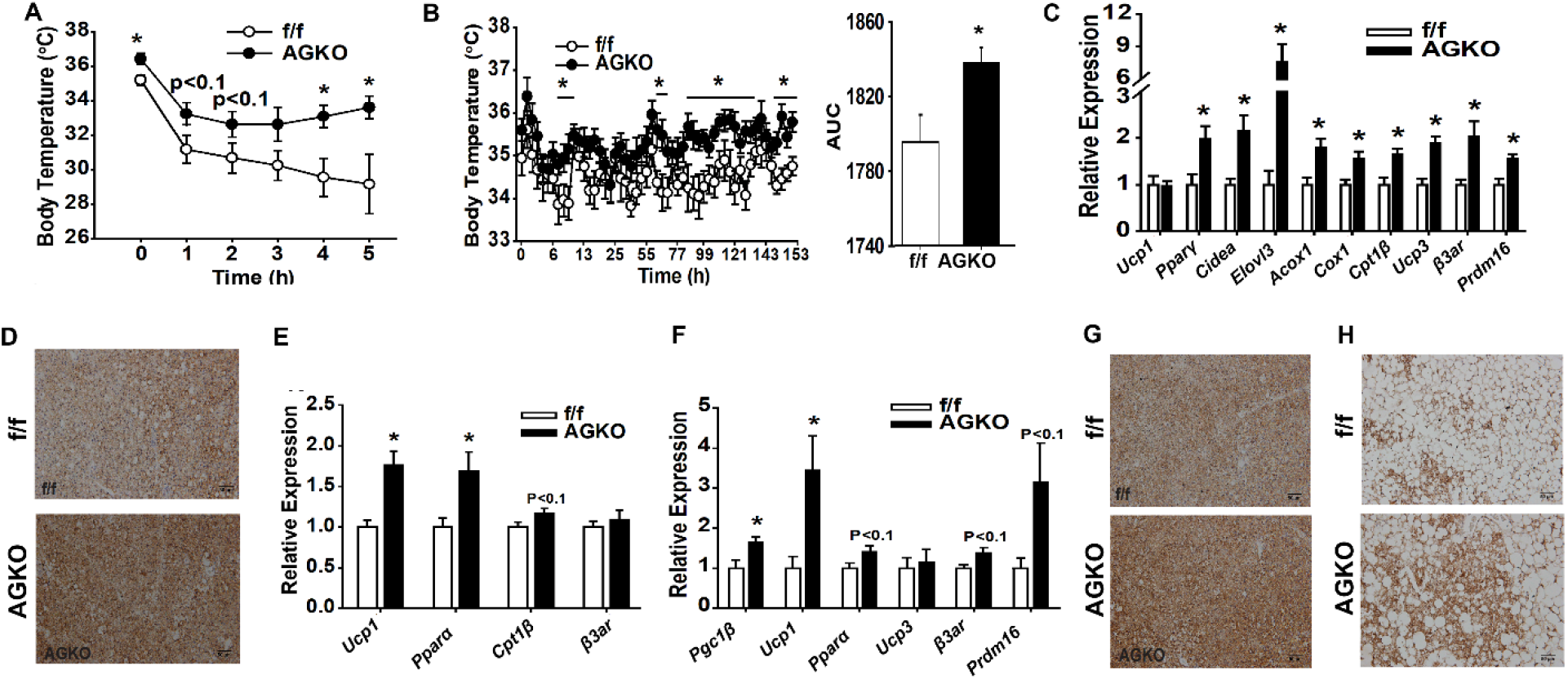
Sensory neuron GHSR knockout mice have increased cold-induced thermogenesis. (A) Acute body temperature measurements in 4°C exposed f/f and AGKO mice; n = 5-7 mice/group. (B) Body temperature (left) and area under the curve (AUC) (right) of body temperature measurements in 7-day 10°C exposed f/f and AGKO mice; n = 10 mice/group. (C) Relative iBAT thermogenic gene expression in 1-day 4°C exposed f/f and AGKO mice; n = 5-7 mice/group. (D) Representative iBAT UCP1 IHC staining in 1-day 4°C exposed f/f and AGKO mice. (E) Relative iBAT thermogenic gene expression in 7-day 10°C exposed f/f and AGKO mice; n = 10 mice/group. (F) Relative iWAT thermogenic gene expression in 7-day 10°C exposed f/f and AGKO mice; n = 10 mice/group. (G) Representative iBAT UCP1 IHC staining in 7-day 10°C exposed f/f and AGKO mice. (H) Representative iWAT UCP1 IHC staining in 7-day 10°C exposed f/f and AGKO mice. Data presented as mean ± SEM. *, p<0.05 versus f/f controls as measured by two-tailed unpaired t-test or two-way ANOVA with Bonferonni post-hoc analysis.

Consistent with their higher cold tolerance, we found AGKO mice had significantly increased thermogenic gene expression and UCP1 IHC staining in iBAT compared to f/f mice during 1-day cold exposure (**Figures 4C-D**). In addition, AGKO mice also had significantly increased thermogenic gene expression in iBAT and iWAT (**Figures 4E-F**), increased UCP1 IHC staining in iBAT (**Figure 4G**) and increased beiging in iWAT (**Figure 4H**) compared to that of f/f mice after the 7-day cold exposure.

### AGKO mice have upregulated sympathetic drive to adipose tissue

Sympathetic drive is necessary for non-shivering thermogenesis (*57*). Indeed, we found that norepinephrine turnover (NETO) in iBAT and iWAT was significantly increased in AGKO versus f/f mice even when housed at room temperature and also in response to a 1-day cold challenge (**Figures 5A-B**), suggesting that AGKO mice have increased sympathetic drive to BAT and WAT. In addition, we also sought to specifically measure sympathetic outflow to adipose tissue by measuring electrophysiological activity of iBAT sympathetic nerves *in vivo,* and found AGKO mice had approximately double the rate of spontaneous sympathetic nerve spikes across 10 minutes (**Figure 5C**). These data support that AGKO mice have increased sympathetic outflow to fat tissues, which is in line with their higher thermogenic capacity and increased energy expenditure during energetic challenges.

**Figure 5.**
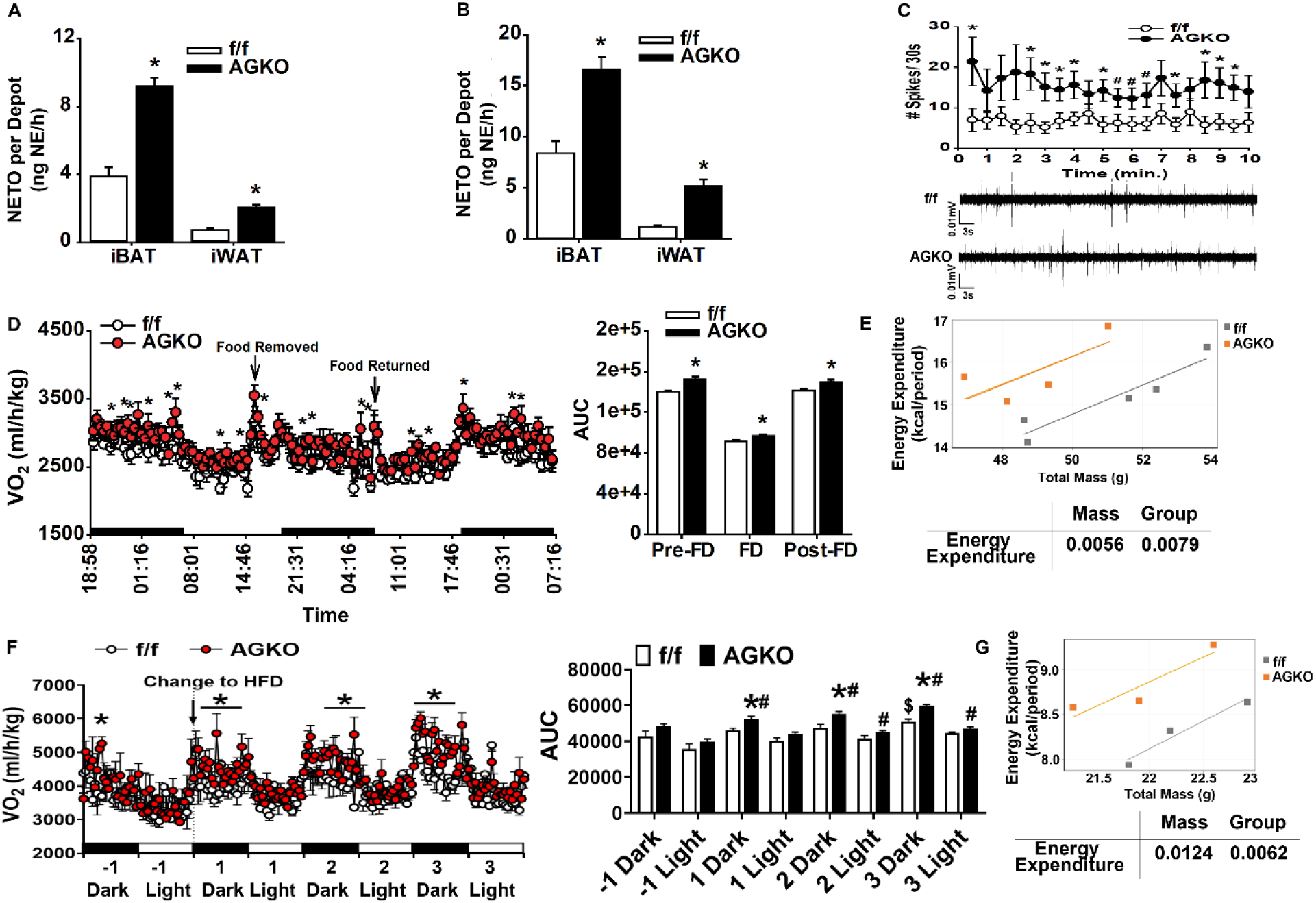
Sensory neuron GHSR knockout mice have increased sympathetic outflow to adipose tissue. (A) iBAT and iWAT norepinephrine turnover (NETO) measurements in room temperature f/f and AGKO mice; n = 9 mice/group. (B) iBAT and iWAT NETO measurements after 1-day 4°C cold exposure in f/f and AGKO mice; n = 9 mice/group. (C) Basal iBAT sympathetic nerve electrophysiology recordings and representative traces in f/f and AGKO mice; n = 9 mice/group. (D) Average oxygen consumption rate (left) and area under the curve (AUC) (right) before, during, and after an overnight food deprivation challenge (Pre-FD, FD and Post-FD, respectively) in f/f and AGKO mice fed HDF for 12 weeks; n = 4-5 mice/group. (E) ANCOVA analysis of energy expenditure data from f/f and AGKO mice during the post-FD phase. n = 4-5 mice/group. (F) Average oxygen consumption rate (left) and area under the curve (AUC) (right) before, and after diet switch from LFD to HFD in 12-week f/f and AGKO mice; n = 3 mice/group. (G) ANCOVA analysis of energy expenditure data from f/f and AGKO mice after the diet switch from LFD to HFD in 12-week f/f and AGKO mice. n = 3 mice/group. Data presented as mean ± SEM. * p<0.05 versus f/f controls, ^#^ p<0.05 vs AGKO -1 Dark or -1 Light, ^$^ p=0.09 vs f/f -1 Dark. Significance was measured by two-tailed unpaired t-test or two-way ANOVA with Bonferonni post-hoc analysis.

We then tested whether sensory GHSR signaling would be important in regulating energy expenditure during energetic challenges, such as food deprivation and in response to HFD. Indeed, we observed that HFD-fed AGKO mice had elevated energy expenditure at baseline before food deprivation (**Figure 5D**), similarly as described in **Figure 3E**. More importantly, AGKO mice also had increased energy expenditure during food deprivation and refeeding phase (**Figure 5D**), while there was no difference in food intake during the refeeding phase (**Figure S9B**) and locomotor activity before, during and after the food deprivation challenge (**Figure S13A**). ANCOVA analysis of energy expenditure during the refeeding phase against body weight suggested a body weight independent effect on the differences in energy expenditure between AGKO and f/f mice (**Figure 5E**). In addition, when switched from LFD to HFD, AGKO mice had significantly increased oxygen consumption starting at 1 day after the diet switch (**Figure 5E**) without any changes in food intake and locomotor activity (**Figure S13B-C**), suggesting a significantly enhanced diet-induced thermogenesis. ANCOVA analysis of energy expenditure against body weight after the HFD switch also suggested a body weight independent effect on the differences in energy expenditure between AGKO and f/f mice during the onset of HFD feeding (**Figure 5G**). Thus, our data further suggest that peripheral sensory GHSR is important in regulating energy expenditure in response to energetic challenges, including food deprivation and HFD challenges.

### Surgical adipose tissue sympathetic denervation abolishes sensory GHSR regulation of energy homeostasis

To further confirm that sensory GHSR regulating body weight and energy homeostasis via adipose tissue sympathetic nervous system, we surgically denervated iBAT and iWAT in AGKO and f/f mice, and challenged them with HFD feeding. As shown in **Figure 6A-B**, sham-operated AGKO mice exhibited higher energy expenditure than that of sham-operated f/f mice during the onset of HFD feeding; however, this increased diet-induced thermogenesis was blunted in iBAT/iWAT denervated AGKO mice (**Figure 6A-B**). ANCOVA analysis of energy expenditure against body weight in sham-operated AGKO and f/f mice after the onset of HFD feeding suggested a body weight independent effect on the differences in energy expenditure between AGKO and f/f mice (**Figure 6C**). In consistence, sham-operated AGKO mice gradually displayed lower body weight than that of f/f mice after the HFD challenge (**Figure 6D-E**); however, this DIO protective effect was abolished in AGKO mice with iBAT and iWAT surgical denervation (**Figure 6D-E**). Sham-operated AGKO mice also showed reduced fat mass compared to that of f/f mice after HFD feeding (**Figure 6F**). Denervation of iBAT and iWAT in AGKO mice also abolished fat mass reduction in AGKO mice compared to that of f/f mice (**Figure 6F**).

**Figure 6.**
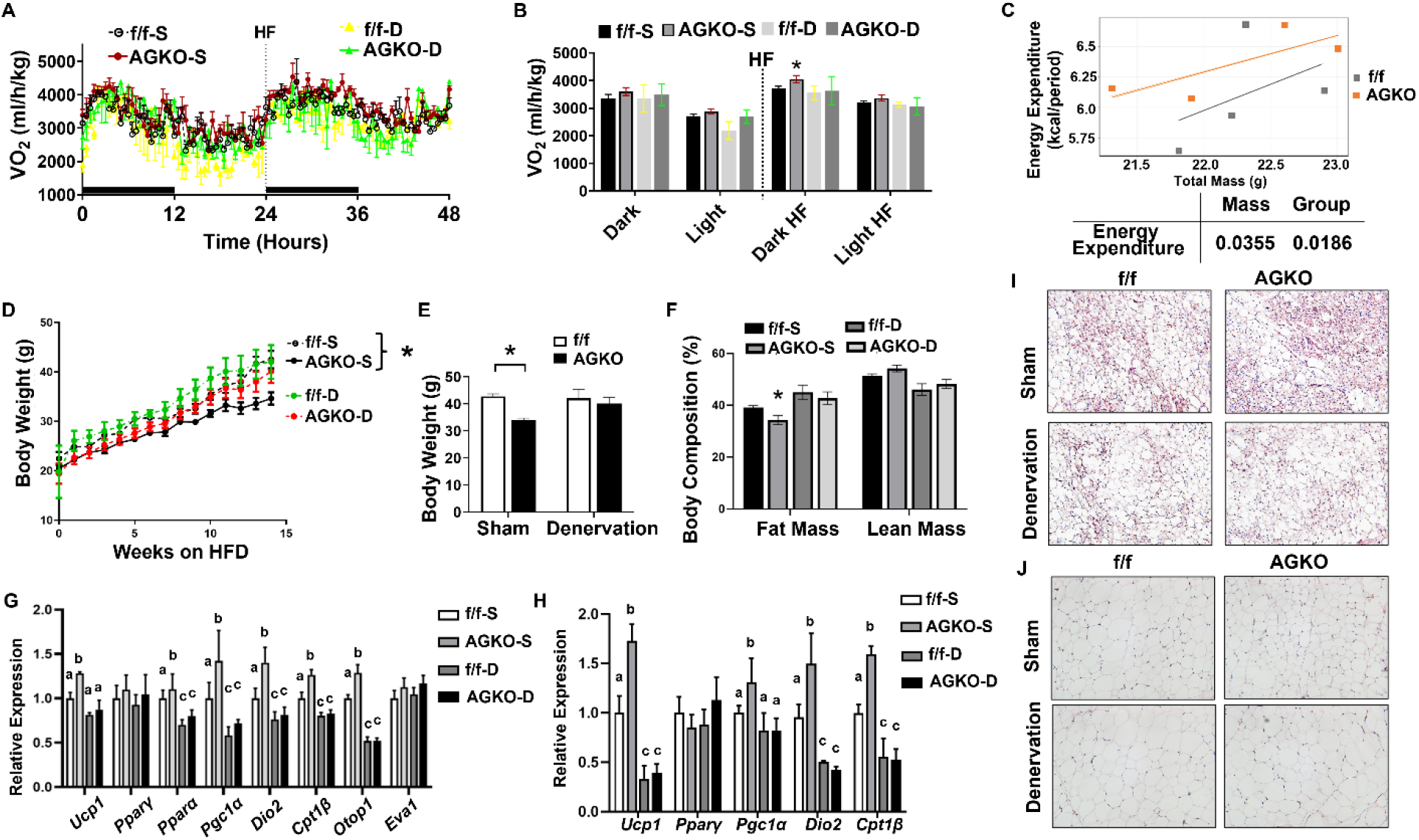
Surgical sympathetic denervation in iBAT and iWAT abolishes the protection from DIO in AGKO mice. (A) Oxygen consumption rate in 8-week-old sham-operated and iBAT/iWAT surgical sympathetic denervated f/f and AGKO mice before and after diet switch from LFD to HFD. n = 4 mice/group. (B) Quantification of oxygen consumption rate in 8-week-old sham-operated and iBAT/iWAT surgical sympathetic denervated f/f and AGKO mice during the dark and light cycle before and after diet switch from LFD to HFD. n = 4 mice/group. (C) ANCOVA analysis of energy expenditure data from sham-operated f/f and AGKO mice after the diet switch from LFD to HFD. n = 4 mice/group. (D) Weekly body mass measurements in sham-operated and iBAT/iWAT denervated f/f and AGKO mice fed 60% HFD; n = 4 mice/group. (E) Body weights in sham-operated and iBAT/iWAT denervated f/f and AGKO mice fed 60% HFD for 14 weeks. n = 4 mice/group. (F) Percentage of fat and lean mass in sham-operated and iBAT/iWAT denervated f/f and AGKO mice fed 60% HFD for 14 weeks. n = 4 mice/group. (G) Gene expression in iBAT of sham-operated and iBAT/iWAT denervated f/f and AGKO mice fed 60% HFD for 14 weeks. n = 4 mice/group. (H) Gene expression in iWAT of sham-operated and iBAT/iWAT denervated f/f and AGKO mice fed 60% HFD for 14 weeks. n = 4 mice/group. (I) Representative H&E staining in iBAT of sham-operated and iBAT/iWAT denervated f/f and AGKO mice fed 60% HFD for 14 weeks. (J) Representative H&E staining in iWAT of sham-operated and iBAT/iWAT denervated f/f and AGKO mice fed 60% HFD for 14 weeks. Data presented as mean ± SEM. * p<0.05 versus f/f controls. In (I) and (J), bars with different lowercase letters are statistically different from each other. Significance was measured by two-tailed unpaired t-test or two-way ANOVA with Bonferonni post-hoc analysis.

Consistent with their increased energy expenditure, iBAT and iWAT from sham-operated AGKO mice exhibited increased expression of thermogenic genes compared to that of f/f mice, including *Ucp1*, *Pparα*, peroxisome proliferative activated receptor gamma coactivator 1 alpha (*Pgc1α*), type 2 deiodinase (*Dio2*), muscle type carnitine palmitoyltransferase 1b (*Cpt1β*) and otopetrin 1 (*Otop1*) in iBAT, and *Ucp1*, *Pgc1α*, *Dio2*, and *Cpt1β* in iWAT (**Figure 6E-F**). Interestingly, we found denervation in f/f mice led to either tended or significant reduction of these thermogenic gene expression, and denervation of iBAT and iWAT in AGKO mice completely blocked the increase in these thermogenic gene expression observed in the sham-operated AGKO mice (**Figure 6G-H**). As a result, denervation of iBAT and iWAT in f/f mice resulted in enlarged adipocyte size in iBAT and iWAT compared to that of sham-operated f/f mice (**Figure 6I-J**). sham-operated AGKO mice showed smaller adipocyte size in iBAT and iWAT when compared to that of sham-operated f/f mice; however, this reduction in adipocyte size was blocked by iBAT and iWAT denervation in AGKO mice (**Figure 6I-J)**.

### There exists extensive crosstalk between non-vagal gastric sensory afferents and iBAT sympathetic outflow

We lastly sought to identify the neuroanatomical basis for the crosstalk between gastric sensory and iBAT sympathetic outflow neurons. We utilized the H129 strain of the anterograde neuronal tract tracer herpes simplex virus 1 (HSV-1 H129) (*58*) to label stomach-projecting sensory neurons and the retrograde neuronal tract tracer pseudorabies virus strain PRV152 (*59, 60*) to label iBAT-projecting sympathetic outflow neurons. To limit infection only to the non-vagal DRG sensory neurons, we performed subdiaphragmatic vagotomy before viral injections. As expected, we observed no H129 and PRV labeling in the nodose ganglia (**Figure S14A-C**). Instead, we observed broad H129, PRV, and double labeled neurons in DRGs (**Figures 7A-B**), sympathetic ganglia (**Figures 7C-D**), and within the spinal cord (**Figures 7E-F**). In DRGs, prominent H129, PRV and double labeling neurons were enriched between T5 and T10 levels (**Figures 7A-B**) that innervate the gut (**Figure S1A**); whereas in sympathetic ganglia, extensive H129, PRV and double labeling neurons were enriched at T1 level (**Figures 7C-D**) that innervates iBAT (*61, 62*). It is suggested that both short feedback and long feedback loops exist between iBAT sympathetic and iWAT sensory circuits (*62*). The short feedback loop consists of neuronal circuitries involving DRGs, spinal cord and sympathetic ganglia; whereas the long feedback loop consists of neuronal circuitries involving DRGs, various brain regions and sympathetic ganglia (*62*). Our data showing broad H129, PRV and double labeling in T5-T10 DRGs innervating the gut as well as in T1 sympathetic ganglia innervating iBAT strongly support an extensive crosstalk between gut sensory and iBAT sympathetic neuronal circuitry, possibly via a similar short feedback loop between the two.

**Figure 7.**
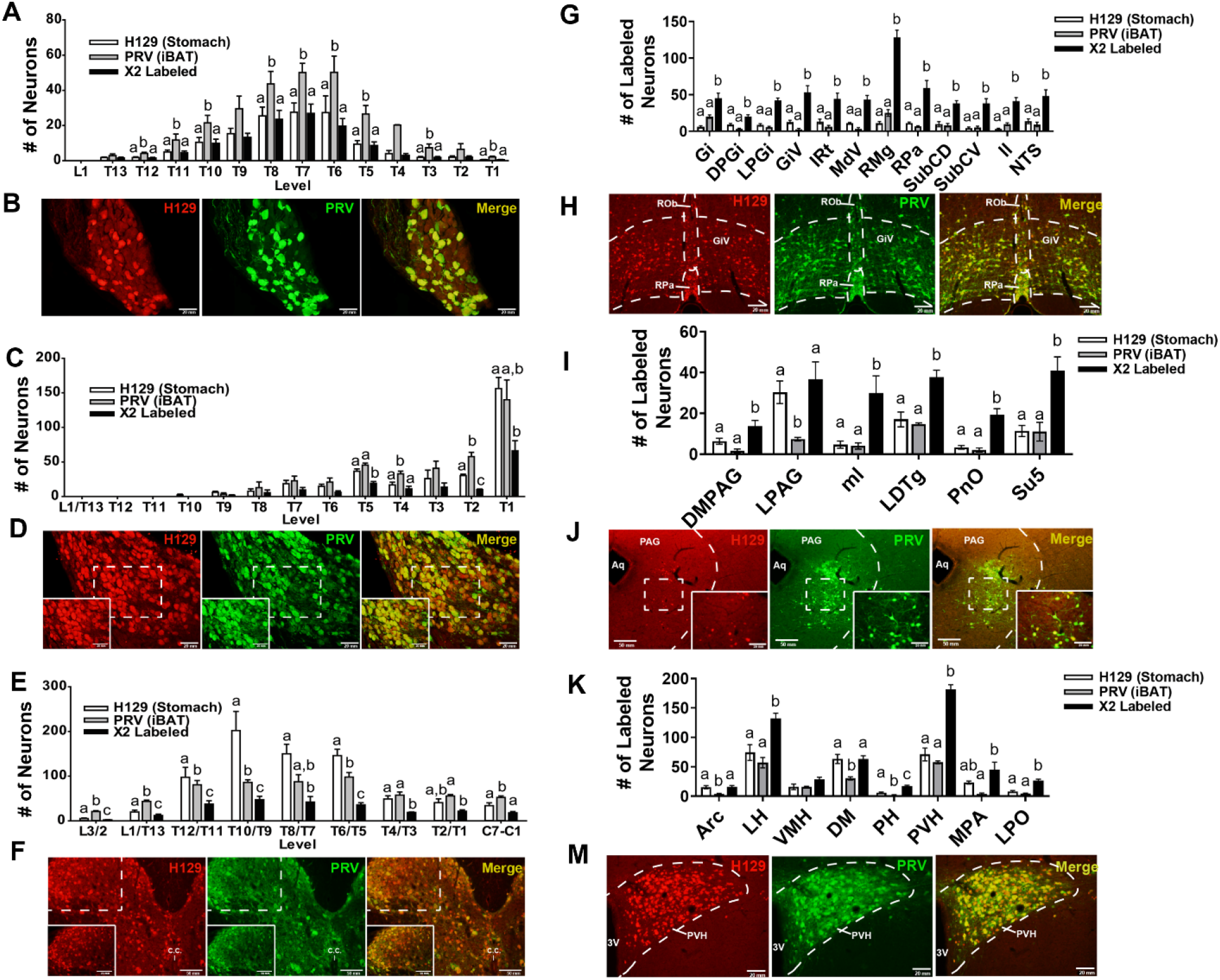
Neuronal crosstalk between stomach-projecting DRG sensory neurons and iBAT-projecting sympathetic outflow neurons. (A) Number of H129-positive, PRV-positive, and double labeled neurons in DRGs; n = 4 mice/group. (B) Representative H129, PRV and double labeling at T10 DRG level. (C) Number of H129-positive, PRV-positive, and double labeled neurons in sympathetic ganglia; n = 4 mice/group. (D) Representative H129, PRV and double labeling at T1 sympathetic ganglia level. (E) Number of H129-positive, PRV-positive, and double labeled neurons in spinal cord sections; n = 4 mice/group. (F) Representative H129, PRV and double labeling at T10/T9 spinal cord level. (G) Number of H129-positive, PRV-positive, and double labeled neurons in hindbrain sections; n = 4 mice/group. (H) Representative image of hindbrain H129, PRV and double labeling (dashed lines denote raphe obscurus (ROb), raphe pallidus (RPa), and gigantocellular reticular nucleus ventral part (GiV)). (I) Number of H129-positive, PRV-positive, and double labeled neurons in midbrain sections; n = 4 mice/group. (J) Representative image of midbrain H129, PRV and double labeling (dashed line denotes periaqueductal gray (PAG); dashed box denotes location of 20x inset). Aq: cerebral aquaduct. (K) Number of H129-positive, PRV-positive, and double labeled neurons in forebrain sections; n = 4 mice/group. (L) Representative image of forebrain H129, PRV and double labeling (dashed line denotes paraventricular hypothalamus (PVH)). 3V: 3^rd^ ventricle. Data presented as mean ± SEM. Bars with the same lowercase letters are not statistically different, and bars with different lowercase letters signify statistical significance (p<0.05) as measured by one-way ANOVA with Bonferonni post-hoc analysis.

In addition, we also found extensive H129, PRV, and double labeling across the forebrain, midbrain, and hindbrain (**Figures 7G-M, S15A-E and Supplemental Table 1**). In hindbrain, neurons with the most prominent double labeling were found in the Doral Raphe Areas, including the raphe magnus nucleus (RMg) and the raphe pallidus nucleus (RPa) (**Figures 7G-H, S15A and Supplemental Table 1**). Other hindbrain areas with extensive double-labeled neurons included nucleus of the solitary tract (NTS), gigantocellular reticular nucleus (Gi), lateral paragigantocellular nucleus (LPGi), gigantocellular reticular nucleus, ventral part (GiV), intermediate reticular nucleus (IRt), medullary reticular nucleus, ventral part (MdV), subcoeruleus nucleus-dorsal/ventral parts (SubCD and SubCV, respectively), lateral lemniscus (ll), parapyramidal nucleus (PPy), rubrospinal tract (rs), and Kolliker-Fuse nucleus (KF) (**Figures 7G- H, S15A-B and Supplemental Table 1**). In midbrain, neurons with high double-labeling were found in periaqueductal gray (PAG) area, laterodorsal tegmental nucleus (LDTg), as well as supratrigeminal nucleus (Su5), pedunculopontine tegmental nucleus (PPTg), medial lemniscus (ml) and pontine reticular nucleus, oral part (PnO) (**Figures 7I-J, S15C and Supplemental Table 1**). In forebrain, neurons with the most abundant double-labeling were found in the paraventricular hypothalamus area (PVH), followed by lateral hypothalamus areas (LH), dorsomedial hypothalamus (DM), and preoptic area, including medial preoptic area (MPA) and lateral preoptic area (LPO) (**Figures 7K-M, S15D-E and Supplemental Table 1**).

## Discussion

Given the multitude of studies demonstrating ghrelin’s role in obesity development (*21, 63, 64*), it is unsurprising ghrelin has emerged as an important regulator of metabolic homeostasis. Previous global and neuronal GHSR knockout studies have shown that ghrelin/GHSR-regulated energy expenditure contributes prominently to obesity resistance in these knockout mice besides the well-known effect of ghrelin/GHSR on food intake (*21, 23, 64*). Here we have identified peripheral sensory GHSR signaling in the regulation of adipose tissue thermogenic capacity and whole-body energy expenditure. We find that GHSR is expressed in DRGs projecting to the gastrointestinal organs, and sensory neuron specific GHSR knockout mice are resistant to DIO. However, this beneficial effect is independent of food intake; instead, it is primarily due to increased energy expenditure and adipose tissue thermogenic capacity, possibly via upregulated sympathetic activity in adipose tissue. To our knowledge this is the first demonstration that sensory neuron GHSR signaling functions to regulate adipose tissue sympathetic outflow and energy expenditure.

While the exact mechanism through which the absence of GHSR signaling increases adipose tissue sympathetic activity necessitates further study, we propose this may be exerted via gut-brain crosstalk through two possible mechanisms. First, ghrelin signaling on sensory neurons may regulate energy expenditure through short feedback loops at the level of the spinal column. In support of this, sensory-sympathetic cross talk at the level of the spinal cord exists in the gastrointestinal system (*65*) as well as between various fat depots (*62, 66, 67*), and spinal column damage impairs normal gastric function despite intact vagal signaling (*68*). Using gastric sensory and iBAT sympathetic viral tracing techniques, we have identified broad sensory-sympathetic colocalization in both sympathetic ganglia and DRGs, and within the spinal column, indicating robust crosstalk across the peripheral nervous system. Second, and more plausibly, GHSR-mediated energy expenditure may be controlled through sensory integration in the brain. Adipose tissue sympathetic outflow neurons are found in numerous hindbrain and forebrain nuclei (*61, 69*), and many of these nuclei concurrently integrate peripheral sensory neuron information to regulate energy expenditure (*66, 70*). Brain receives inputs from temperature-and energy-sensing neurons through hypothalamic and brain stem areas including the preoptic area (POA), ARC hypothalamus, and NTS, and sends efferent fibers to sympathetic premotor neurons in the rostral medullary raphe (RMR), including RPa and RMg nucleus, or project directly to spinal preganglionic neurons (*71, 72*). Our sensory-sympathetic viral tracing analysis showing broad sensory-sympathetic colocalization in these similar hindbrain, midbrain, and forebrain nuclei further emphasizes the importance of peripheral sensory GHSR signaling in integrating gut sensory input into a neuronal network that regulates sympathetic outflow to adipose tissue, which in turn regulates whole body energy homeostasis.

GHSR is also expressed in vagal afferent neurons (VAN) located in the nodose ganglia (*29*) and vagal GHSRs were initially hypothesized to be necessary and sufficient for ghrelin signaling (*30*). Yet subsequent studies in vagotomized animals found no change in ghrelin-induced food intake and adaptation responses to cold (*33, 34*). In addition, a recent study showed that, while adeno-associated virus (AAV)-mediated *Ghsr* knockdown in the nodose ganglia in rats blunted ghrelin-induced food intake in rats (*29*), nodose ganglia *Ghsr* knockdown did not affect 24-hour food intake; further, unlike the whole body or neuronal *Ghsr* deficient mice that exhibited DIO resistance (*21, 23*), nodose ganglia *Ghsr* knockdown increased meal frequency, delayed gastric emptying and resulted in a slightly higher body weight over time in rats fed a regular chow diet (*29*). However, diet-induced obesity was not studied (*29*), and thus, further studies may be needed to delineate the role of VAN GHSR in regulating energy homeostasis. On the other hand, our findings here demonstrate that *Ghsr* is expressed in DRGs, and that sensory neuron *Ghsr* deletion imparts a beneficial metabolic phenotype. Thus, our data suggest a compensatory mechanism involving DRG *Ghsr*-expressing sensory neuronal circuits that may be sufficient to mediate ghrelin’s peripheral effects independent of the vagus.

Selective GHSR restoration in AgRP neurons or hindbrain *Phox2b* expressing neurons restores fasting glucose levels suggesting these neurons are sufficient to mediate ghrelin’s glucoregulatory effects (*28, 50*). Since Phox2b is expressed in both hindbrain and peripheral sensory neurons (*73, 74*), it is therefore difficult to delineate the central or peripheral mechanism mediating this effect. Our results demonstrate that peripheral sensory neuron specific GHSR knockout has no effect on fasting glucose levels in chow-fed animals, which would suggest that the improved glucoregulation in Phox2b GHSR-restored mice involves a central mechanism likely in hindbrain nuclei. On the other hand, the improvement of insulin sensitivity in HFD-fed AGKO mice may be secondary to their protection from diet-induced obesity.

In sum, our results demonstrate a novel peripheral ghrelin signaling pathway that responds to energetic challenges and regulates metabolic homeostasis. As this pathway is independent of central ghrelin signaling, it strongly suggests the presence of multiple ghrelin-involved mechanisms that regulate whole body energy homeostasis. Hence, peripheral ghrelin signal intervention may provide novel therapeutic approaches that increase basal energy expenditure and prevent or reverse obesity.

## Acknowledgements

We thank Dr. John Garretson for helpful technical inputs for the electrophysiology recording of iBAT sympathetic nerve efferent activity experiment, Dr. Lynn Enquist for providing the H129 and PRV152 viruses (Virus Center Grant P40RR018604) and helpful technical inputs in the viral tracing experiments; and Dr. Robert C. Ritter for helpful technical inputs in the vagotomy and gastrointestinal organ microinjection experiments. This research was supported by NIH R01DK125081, R01DK118106, and American Diabetes Association (ADA) 1-18-IBS-260 to B.X.; NIH R01DK118106, R01DK115740, and ADA 1-18-IBS-348 to H.S.; NIH DK116425 to M.A.T., and R01AG064869 to Y.S.

## Author contribution

MAT participated in conceiving and designing the study, performed most of the experiments and data analysis, and wrote the first draft of the manuscript; XC performed vagotomy and viral tracing experiments and analyzed data; LA performed electrophysiology recording in DRG neurons; LA and FS helped MAT with the calcium measurement; QC and JJ assisted in various experiments; JZ provided technical inputs and review/edits of the manuscript; YS provided *Syn1-Cre;Ghsr^f/f^* mice, technical inputs and review/edits of the manuscript; BX and HS conceived and designed study, interpreted the data, and wrote and finalized the manuscript.

## Data Availability

The data supporting the findings of this study are available from the corresponding author upon reasonable request.

## Methods

### Animals

Adult C57BL/6J mice were obtained from the Jackson Laboratory (Stock No: 000664). Adult *Ghsr*-IRES-tauGFP mice were obtained from the Jackson Laboratory (Stock No: 019908)(*40*). To generate a sensory neuron specific Ghsr knockout model, *Syn1-Cre;Ghsr^f/f^* mice (*23*) were first bred with wild type C57BL/6J mice to obtain *Ghsr^f/f^* mice, which were then bred with *Advillin-Cre* mice (Kindly provided by Dr. Fan Wang from Duke University) (*49*) to generate *Advillin-Cre*^+/-^;*Ghsr^f/f^* mice (henceforth referred to as AGKO). Male AGKO or f/f littermate mice were fed either a standard chow diet (catalog no.: 5001; LabDiet; Purina, St. Louis, MO) or high fat diet (HFD) consisting of 60% calories from fat (catalog no. D12492; Research Diets, New Brunswick, NJ). In some of the experiments, a low-fat diet (LFD) (Research Diets D12450B, 10% calories from fat) was used as a control diet. Body mass measurements were taken weekly for the duration of the experiment. Animals were maintained at ∼23°C with 12-hour light/dark cycles (0700-1900 light). All procedures were approved by the Georgia State University Institutional Animal Care and Use Committee and were in compliance with the Public Health Service and United States Department of Agriculture guidelines.

### Physiology Measurements

Glucose and insulin tolerance tests were performed on male AGKO or f/f littermate mice after 12 weeks on either Chow or high-fat diet (HFD). For glucose tolerance tests, animals were fasted for 16 hours overnight. Baseline glucose measurements were obtained from a tail nick using a OneTouch Ultra Blood Glucose Meter and glucose test strips (LifeScan, Inc., Milpitas, CA). Following the initial measurement, mice were intraperitoneally injected with a 20% dextrose solution (1g/kg body mass), and subsequent blood glucose measurements were performed at 15, 30, 60, 90, and 120 minutes after the glucose injection. For the insulin tolerance test (ITT), mice were fasted for 4 hours prior to the start of the procedure, and baseline blood glucose was measured from a small tail nick. Mice were then intraperitoneally injected with 1.0 U/kg insulin (Humulin R; Eli Lilly, Indianapolis, IN), and blood glucose was measured at the same time points used for the GTT. Energy expenditure, oxygen consumption, carbon dioxide production, respiratory exchange ratio, and physical activity data were collected for male AGKO or f/f littermate mice using a TSE PhenoMaster metabolic chamber system (TSE Systems, Chesterfield, MO) after 12 weeks on Chow or HFD. Animals were allowed a 48-hour acclimation period prior to the start of data collection and then returned to their home cage at the end of the experiment.

### Body Composition Measurements

Body composition measurements were taken for HFD-fed male AGKO or f/f littermate mice using our Minispec LF90 TD-NMR analyzer (Bruker Optics). For all measurements, animals were briefly removed from their home cage, placed in the body composition analyzer, and then immediately returned to their home cage. Body composition measurements were taken for all animals prior to the start of HFD feeding, after 8 weeks of HFD feeding, and then again at the conclusion of the experiment.

### Tissue Collection

At the end of each experiment, mice were euthanized using carbon dioxide inhalation. Blood serum was collected and stored at −80°C until analysis. Animals were then perfused with ice cold 0.1M PBS and fat pads (brown (iBAT), epididymal (eWAT), and inguinal (iWAT)), liver, brain, and DRGs (L5-T5 levels) dissected and flash frozen in liquid nitrogen. For brain microdissection, the brain was placed on an ice cold cutting block and 1 mm sagittal sections taken from both sides of the midline. The arcuate hypothalamus, paraventricular hypothalamus, ventromedial hypothalamus, and ventral tegmental area were then microdissected according to a mouse brain atlas (Paxinos and Franklin (*75*)) and flash frozen in liquid nitrogen. A sample of each fat pad and liver was taken prior to freezing and placed in formalin for later histological analysis. In a separate set of *Ghsr-GFP* mice used for immunohistochemical analysis, animals were euthanized by carbon dioxide inhalation and then transcardially perfused with 75 ml 0.9% heparinized saline followed by 150 ml 4.0% paraformaldehyde in 0.1 M PBS in sterile filtered dH2O. DRGs (L5-T5 levels) were subsequently removed and dehydrated in an 18% sucrose solution until immunohistochemical analysis.

### Fast Blue Injections

1% Fast Blue in sterile dH_2_O (17740-1, Polysciences, Warrington, PA) was used as a fluorescent retrograde neuronal tracer to identify gastrointestinal-projecting sensory neurons using a modified method as previously described (*76*). In brief, animals were deeply anesthetized with 1-2% isoflurane, and a small incision was made on the ventral midline to expose the peritoneal cavity and discrete gut regions (i.e. stomach and small and large intestine). A series of 0.5-1 µl injections were given across 4 loci along greater curvature of the stomach or across 8 evenly spaced loci along the small or large intestine through a 26-gauge needle and syringe. The peritoneal cavity was then closed with sterile dissolvable sutures and the skin incision closed with sterile wound clips. Animals were given 7 days to recover and allow for retrograde Fast Blue transport before euthanization and tissue extraction as described above.

### Subdiaphragmatic Vagotomy and Viral Injections

All viral injections were performed according to Biosafety Level 2 standards. 4 adult (2-3 mo.) male C57BL/6 mice were initially single housed in ACS cages at room temperature. For subdiaphragmatic vagotomy, animals were anesthetized with 1-2% isoflurane and a small incision made on the ventral midline to isolate the stomach. The ventral vagus nerve is isolated where it exits the diaphragm, the gastric branches identified, and a small 1-2 mm section of each branch excised. Following the vagotomy, the stomach received modified H129 strain of herpes simplex virus-1 (gift from Dr. Lynn Enquist, Princeton University, Princeton, NJ; Virus Center Grant P40RR018604) (HSV-1 H129) (*58*) microinjections at 5 loci along greater curvature of the stomach (5.12 × 10^8^ pfu/ml; 150 nl/locus), and the needle left in place for 1 minute following each injection to prevent efflux. The peritoneal cavity was then closed with sterile dissolvable sutures and the skin incision closed with sterile wound clips.

While still under anesthesia, mice were flipped over on their stomach, and a small skin incision made above iBAT. Microinjections of modified pseudorabies virus PRV152 containing cytomegalovirus-enhanced green fluorescent protein (CMV-EGFP) reporter gene cassette inserted into the gG locus of the viral genome (gift from Dr. Lynn Enquist, Princeton University, Princeton, NJ; Virus Center Grant P40RR018604) (*59, 60*) were given across five evenly distributed loci (2 × 10^9^ pfu/ml, 150 nl/locus) for each iBAT pad (10 total injections per animal) and the needle left in place for 1 minute following each injection to prevent efflux. After the final injection, the skin was closed with sterile wound clip staples and the animal returned to its cage. Animals were kept isolated in their home cage for 6 days to recover and allow for viral transport before euthanization and tissue extraction as described above. To ensure the specificity of viral injection, we have previously tested PRV and HSV-1 H129 injections for potential viral leakage by placing the virus on the surface of the tissue and found no positive labeling in the DRGs, SNS ganglia, spinal cord, and brain (*61, 62*).

### Histological Analysis

For DRG GFP immunohistochemical quantification in our *Ghsr-GFP* mice injected with Fast Blue, DRGs (L5-T5 levels) were removed from the sucrose solution and sliced in a cryostat at 25 µm. DRG sections were taken in series across three slides (SuperFrost Plus slides (Thermo Fisher Scientific); 8 sections/slide total) to prevent a single neuron being present multiple times on the same slide as previously described (*66, 76*). For immunohistochemical staining, in brief, slides were washed 3X15 min in sterile 0.1M PBS on a rotator. DRG sections were blocked with 5% normal horse serum containing 0.3% Triton X-100 in 0.1M PBS for 1 h. Slides were then incubated overnight at room temperature with rabbit anti-cFOS (1:500; sc-52, Santa Cruz Biotechnology, Santa Cruz, CA) and chicken anti-GFP (1:400; GFP2020, AvesLabs, Tigard, OR) antibodies in 0.1M PBS with 0.3% Triton X-100 and 10% normal horse serum. Next, slides were washed 3X 15 min in 0.1M PBS and then incubated at 4°C for 24 h with donkey anti-chicken-488 (1:1000; 703-545-155, Jackson Immunoresearch, West Grove, PA) and donkey anti-rabbit-CY3 (1:400; 711-165-152, Jackson Immunoresearch, West Grove, PA) in 0.1M PBS. Slides were then washed 3 × 15 min with 0.1M PBS and coverslipped with Prolong antifade fluorescent medium (Life Technologies, Grand Island, NY). Images were captured using an Olympus DP73 photomicroscope and CellSens software (Olympus, Waltham, MA). For GFP, cFOS, and Fast Blue quantification, total cell numbers were estimated by counting the number of fluorescently positive neurons on a single section and then multiplying this number by 8 as each slide contains 8 sections as previously described (*76*).

Immunohistochemical analysis of DRGs, sympathetic ganglia, spinal cords and brains from PRV152 and HSV-1 H129-injected animals was performed according to previous descriptions (*61, 62*). Briefly, animals were euthanized and transcardially perfused with 0.9% heparinized saline followed by 4% paraformaldehyde. The DRGs, sympathetic ganglia, spinal cords and brains were removed and placed in the same fixative overnight and then immersed in a cryoprotectant solution of 18% sucrose (DRGs and sympathetic ganglia) or 30% sucrose (brains and spinal cords). The brains were sectioned at 30μm on a freezing-stage sliding microtome, and spinal cords and ganglia were sectioned at 40 and 16 µm, respectively, on a Leica cryostat and directly transferred to slides (Superfrost Plus; VWR International, Arlington, IL). Sections of brains, spinal cords, and ganglia were stored in 0.1 M PBS with 0.1% sodium azide (NaN3) at 4°C until double fluorescent immunohistochemistry (IHC) was performed. Every 4th section of the ganglia, spinal cords and brains was processed for double-fluorescent immunohistochemistry (IHC) against HSV-1 and GFP. For immunohistochemistry, free-floating brain sections were subsequently rinsed in 0.1 M PBS (2X15 min) and 0.1% sodium borohydride in 0.1 M PBS to reduce autofluorescence, followed by 30 min incubation in a blocking solution of 10% normal donkey serum (NDS) and 0.4% Triton X-100 in 0.1 M PBS. Sections were then incubated in the mixture of primary rabbit anti-HSV-1, (1:2000; B0114, DakoCytomation, Carpinteria, CA) and chicken anti-GFP (1:400; GFP2020, AvesLabs, Tigard, OR) antibodies with 2% NDS in 0.1 M PBS for 18 h. Subsequently, sections were incubated in the appropriate mixture of the secondary donkey anti-rabbit Cy3 (1:700; 711-165-152, Jackson Immunoresearch, West Grove, PA) and donkey anti-chicken-488 (1:1000; 703-545-155, Jackson Immunoresearch, West Grove, PA) antibodies in 0.1 M PBS for 2 h. Immunohistochemistry for HSV-1 and PRV152 in DRG, sympathetic ganglia and spinal cords was conducted directly on the slides using primary rabbit anti-HSV-1 (1:100; B0114, DakoCytomation, Carpinteria, CA) and chicken anti-GFP (1:400; GFP2020, AvesLabs, Tigard, OR) antibodies. All steps were performed at room temperature. Sections were mounted onto slides and cover slipped using ProLong Gold Antifade Reagent (Life Technologies, Grand Island, NY). H129 and PRV-labeled neurons were considered positive based on cell size, shape, and fluorescent intensity. Images were captured using an Olympus DP73 photomicroscope and were adjusted for brightness, contrast, sharpness, and overlaying of double-labeled neurons using Adobe Photoshop CS5 (Adobe Systems, San Jose, CA). Absolute values of single-or double-labeled neurons were counted across each nucleus/region within each mouse and quantified using the manual tag feature of Adobe Photoshop thus ensuring each neuron was counted once. A mouse brain atlas was used as a reference to identify brain sites (*77*).

For adipose tissue immunohistochemical analysis, samples were removed from formalin, dehydrated through a series of increasing isopropanol washes, and embedded in paraffin blocks. iBAT, iWAT, and eWAT were sliced at 5 µm on a rotating microtome and mounted on SuperFrost Plus slides (Thermo Fisher Scientific). Slides were deparaffinized and hydrated by washing in p-xylenes followed by a series of decreasing ethanol washes. Sections were next incubated in DAKO antigen retrieval solution (DAKO, S1699) and heated in a microwave for 14 minutes total. Rabbit anti-UCP1 antibody (1:150; ab10983, Abcam) was applied to all sections and then incubated overnight at 4°C. Slides were then washed 4 × 30s in a tris buffer solution and the secondary biotinylated mouse adsorbed donkey anti-rabbit antibody (Jackson ImmunoResearch, 711-065-152) applied to all sections and incubated for 30 min. at room temperature. Slides were washed 4 × 30s in tris buffer and then incubated in an ABC solution (Vector Labs, PK-6100) for 30 minutes at room temperature. Sections were then washed 2 × 30s in 1M tris-HCl solution and then incubated in a DAB reaction solution (Vector Labs, SK-4100) for 5-15 min. Slides were washed in dH2O for 5 min. and then dehydrated through graded alcohol washes followed by a final p-xylene wash. Slides were then coverslipped with permount (Fisher, SP15-100), allowed to dry, and imaged with an Olympus DP73 photomicroscope and CellSens software (Olympus, Waltham, MA).

For hematoxylin and eosin adipose tissue histology, samples were removed from formalin, dehydrated through a series of increasing isopropanol washes, and embedded in paraffin blocks. iBAT, iWAT, and eWAT were sliced at 5 µm on a rotating microtome and mounted on SuperFrost Plus slides (Thermo Fisher Scientific). Tissue morphology was visually analyzed using hematoxylin and eosin staining (Sigma-Aldrich). Adipose histology images were captured using an Olympus DP73 photomicroscope and CellSens software (Olympus, Waltham, MA).

### In Vitro DRG Culture

Primary DRG neurons were isolated and cultured as described previously(*78*). Briefly, adult *Ghsr-GFP* mice were euthanized by carbon dioxide inhalation, and DRGs (L5-T5 levels) immediately removed and placed in ice cold 0.1M PBS with 20mM HEPES. This solution was then replaced with 1mg/ml collagenase 1 (Worthington Biochemical Corporation CLS-1) in 0.1M PBS and 20mM HEPES for 60-90 min at 37°C followed by 0.05% Trypsin-EDTA (Sigma T9201) solution in 0.1M PBS and 20mM HEPES for 7 min at 37°C. DRGs were then washed 2X with 1ml complete neurobasal media (Fisher 10888022) with 1X B-27 (Fisher 17504044) and 200µM L-glutamine (Fisher 25030149). Next, 600µl complete neurobasal media was added and the DRGs triturated 30X. Dissociated neurons were filtered through a 100µm cell strainer into a new microcentrifuge tube and then washed 2X with 500µl complete neurobasal media. Neurons were then centrifuged for 5 min at 200g, the supernatant removed and replaced with 1ml complete neurobasal media, triturated 3X, and then seeded onto Poly-D-Lysine and Laminin coated coverslips. Cultured neurons were transferred to a 37°C + 0.5% CO_2_ incubator until use.

### *In vitro* Calcium Imaging and *in vitro* whole-cell path-clamp electrophysiology

For *in vitro* calcium imaging, an Olympus microscope (BX51 W1F, Olympus, Japan) equipped with water-immersive objective (40X), ratiometric dichroic mirror with 340/380 split, 400 nm long-pass emission filter was used for acquiring ratiometric time-lapse images of Fura-2 fluorescence in DRG neurons as described previously (*79-84*). In brief, coverslips containing cultured DRG neurons were transferred to a new 35mm dish containing 5µM Fura-2- acetoxymethyl ester (Fura-2 AM) (Millipore Sigma 344905) in complete neurobasal media and placed in a 37°C + 0.5% CO2 incubator for 30 min. Neurons were then washed 3X with an extracellular recording solution composed of 152mM NaCl, 2.8mM KCl, 10mM HEPES, 2mM CaCl2, and 10mM glucose (pH = 7.3) and placed in a 37°C for additional 20-30 min to allow for complete de-esterification of Fura-2 AM. GFP-positive neurons were identified using GFP excitation and emission filters. Imaging data acquisition and analysis were performed using Live Acquisition and Analysis software (Till Photonics, FEI, Germany). Time-lapse ratiometric images were acquired every 30 seconds with 100 ms exposure at 340 nm and 380 nm excitation wavelengths and 400 nm long-path emission filter. The ratio of the emission fluorescence intensities at 340nm excitation and 380nm excitation wavelengths was used to assess changes in intracellular calcium concentration. During the experiment, baseline fluorescence was measured for 5-10 min. Next, either 100nM ghrelin (Bachem 4033076) or vehicle (extracellular recording solution) was administered and the calcium ratiometric images were measured for additional 30-60 min. At the end of each recording, a depolarization by 60mM KCl was given as a positive control for neuronal viability, and only neurons that responded to KCl treatment with an increased ratio of fluorescent intensity of 340nm over 380nm by KCl were used for analysis. The change in intracellular calcium concentration was indicated as ΔF/F_b_, which was calculated by first taking the average baseline fluorescence intensity ratio of 340nm/380nm across the initial 5- 10 minutes of the study (F_b_). Next, change in fluorescence ratio was calculated at each timepoint X using the formula ΔF/F_b_=[(F_x_ -F_b_)/F_b_].

For electrophysiological recordings, DRG neurons were extracted from *Ghsr-GFP* mice and dissociated as described above. Dissociated neurons were plated into 35 mm plastic dishes coated with poly-D-lysine and used for recording 3-5 days after plating. GFP-labeled DRG neurons were visualized using a GFP-filtered excitation and emission fluorescent light under an inverted microscope. The external solution (pH 7.3) consisted of 152 mM NaCl, 2.8 mM KCL, 10 mM HEPES, 2 mM CaCl2, and 10 mM glucose. The intracellular solution (pH 7.3) consisted of 130 mM K-gluconate, 10 mM HELES, 0.6 mM EGTA, 0.3 mM K-ATP, 0.3 mM Na-GTP, and 10 mM phosphocreatine. The whole-cell patch-clamp recording acquisitions and data analysis were performed using a Multiclamp 700B amplifier, Digidata 1440A interphase, and pClamp 10 software (Molecular Devices, Union City, CA) under current clamp mode as we described previously (*85-87*).

### Electrophysiological Recordings of iBAT Sympathetic Nerve Efferent Activity

Electrophysiological recording of iBAT sympathetic nerve afferent activity was conducted using a modified method as previously described (*76*). Briefly, 9 f/f and 9 AGKO mice were anesthetized with 1-2% isoflurane and a small interscapular incision made. iBAT was isolated and connective tissue removed to allow the entire fat pad to be flipped and tucked into the skin incision to expose the sympathetic nerves. A single nerve was isolated, severed, and then placed over 32-gauge silver hook electrodes to measure efferent activity. Mineral oil was immediately applied to cover the nerve and electrodes and applied periodically throughout the recording to ensure the nerve was always covered and prevent the drying of tissues. Animals were maintained under 1-2% isoflurane anesthesia for the entirety of the 10 minutes recording session, and then immediately euthanized after the recording was terminated.

For all recordings, a differential AC amplifier (Model 1700, A-M Systems) was used to amplify extracellular signals 10K times with a 100 Hz low pass filter and 1000 Hz high pass filter. Electrophysiological activity was digitized with Digidata 1440a data acquisition system (1440a; Molecular Devices), and recorded with Clampex 10.3 software at a 20,000 Hz sampling rate. Spike number was analyzed within Clampfit 10.2 software based on a voltage threshold at least two standard deviations above background, non-signal noise. All identified waveforms were visually analyzed, and any aberrant non-action potential spikes erroneously counted by the software were removed from the analysis.

### Quantitative Gene and Protein Analysis

RNA from frozen tissues was extracted using TRIzol Reagent (Thermo Fisher) according to the manufacturer’s instructions. Gene expression was measured using a 7500-Fast RT-PCR machine (Applied Biosystems), ABI Universal PCR Master Mix (Applied Biosystems, Foster City, CA), and primer and probe sets purchased from Applied BioSystems. Relative gene expression was determined using the ΔΔCt method with cyclophilin as an internal control (*88*).

### Norepinephrine Turnover

Norepinephrine (NE) turnover (NETO) was measured as we described previously (*61, 89*). Briefly, for room temperature NETO measurements, 18 AGKO and 18 f/f mice were single housed in shoebox cages and handled daily for one week prior to the start of the experiment to adapt them to handling and reduce stress-induced NE release. At the start of the experiment, 9 f/f and 9 AGKO mice were injected with saline, and 9 f/f and 9AGKO mice were injected with AMPT (300mg/kg I.P.) (Sigma-Aldrich M3281-1g), an active competitive inhibitor for tyrosine hydroxylase (TH), the rate limiting enzyme for NE production. Two hours later, saline or 150mg/kg IP AMPT was injected again. Animals were then euthanized two hours after the second saline/AMPT injection (4h total from start of experiment) and tissues extracted and flash frozen.

For 1-day 4°C cold exposure NETO measurements, 18 AGKO and 18 f/f mice were single housed in shoebox cages and handled daily exactly as described above. At the start of the experiment, animals were housed in new shoebox cages and the cotton nestlet removed. After 20 hours at 5°C, 9 f/f and 9 AGKO mice were injected with saline, and 9 f/f and 9AGKO mice were injected with 300mg/kg AMPT I.P. (Sigma-Aldrich M3281-1g). Two hours later, saline or 150mg/kg IP AMPT was injected again. Animals were then euthanized two hours after the second saline/AMPT injection (4h total from start of experiment) and tissues extracted and flash frozen.

For both experiments, iBAT, eWAT, and iWAT were processed and NE extracted with dihydroxybenzylamine (Sigma-Aldrich, St. Louis, MO), an internal control for extraction efficiency. NETO measurements were done as described previously (*90*). NETO was calculated using the following formula: k = (lg[NE]0 − lg[NE]4)/(0.434 × 4) and K = k[NE]0, where k is the constant rate of NE efflux, [NE]0 is the initial NE concentration, [NE]4 is the final NE concentration, and K = NETO.

### Cold Exposure

Two weeks prior to the start of cold exposure, 10-week-old AGKO and f/f littermate mice were implanted with temperature transponders (IPTT-300, BMDS, Seaford, DE). In brief, animals were deeply anesthetized with 1-2% isoflurane and a small incision made along the ventral midline to expose the peritoneal cavity. Sterile temperature transponders were inserted into the peritoneal cavity, the incision closed with sterile dissolvable sutures, and the skin incision closed with sterile wound clips. Following transponder implantation, animals were singly housed in shoebox cages with only corn cob bedding for recovery. At the start of the experiment, baseline body temperature measurements were taken for all animals at room temperature before the cold challenge. Core body temperature measurements were taken for all mice every hour for the first 8 hours of the study, and then every 2 hours through 24 h. Measurements were taken every 2 hours from 0700 to 1900 for the remaining 6 days. At the end of the experiment, animals were removed from the cooler, immediately euthanized, and tissues extracted as described above. Mice were either exposed to acute cold (4°C) for 1 day or chronic cold (10°C) for 7 days.

### Food Intake Studies

For ghrelin-induced food intake measurements, single housed ad libitum Chow-fed or HFD-fed AGKO and f/f littermate mice were given a 0.5 mg/kg subcutaneous ghrelin challenge (4033076, Bachem, Torrance, CA) at 0800. Food intake was measured at 30, 60, 90, and 120 min post-injection. For food deprivation-induced food intake studies, single housed Chow-fed or HFD-fed AGKO and f/f littermate mice were food deprived for 16 hours. At 0800, food was returned and food intake measured at 30, 60, 90, and 120 min post-refeeding.

### Plasma ghrelin measurement

For plasma acyl-ghrelin measurement, blood was collected into ice-cold EDTA-coated Eppendorf tubes, followed by addition of P-hydroxymercuribenzoic acid (final concentration 1 mmol/L; Sigma-Aldrich, St. Louis, MO), centrifugation at 4°C at 1,500 g x 15 min to separate plasma, addition of 1N HCl to achieve a final concentration of 0.1N, and storage at -80°C prior to processing for plasma acyl-ghrelin measurement. Acyl-ghrelin concentrations in mouse plasma samples were determined by commercial ELISA kits (EMD Millipore, Billerica, MD; Catalog #EZRGRA-90K) (*91*).

### Statistical Analysis

All data were expressed as mean ± S.E. Experimental data were analyzed by a Student’s t-test, one-way or two-way ANOVA followed by Bonferonni post-hoc analysis where appropriate. For all experiments, differences among groups was considered statistically significant at P<0.05 and trending toward significance at P≤0.1.

## References

1. J. O. Hill, H. R. Wyatt, J. C. Peters, Energy balance and obesity. Circulation 126, 126–132 (2012).

2. G. J. Dockray, Gastrointestinal hormones and the dialogue between gut and brain. J Physiol 592, 2927–2941 (2014).

3. T. H. Moran, Gut peptides in the control of food intake. Int.J Obes.(Lond) 33 **Suppl 1**, S7–10 (2009).

4. C. Blouet, G. J. Schwartz, Duodenal lipid sensing activates vagal afferents to regulate non-shivering brown fat thermogenesis in rats. PLoS One 7, e51898 (2012).

5. M. Kojima et al., Ghrelin is a growth-hormone-releasing acylated peptide from stomach. Nature 402, 656–660 (1999).

6. F. Broglio et al., Ghrelin, a natural GH secretagogue produced by the stomach, induces hyperglycemia and reduces insulin secretion in humans. J Clin Endocrinol Metab 86, 5083–5086 (2001).

7. K. Fujino et al., Ghrelin induces fasted motor activity of the gastrointestinal tract in conscious fed rats. J Physiol 550, 227–240 (2003).

8. M. S. Huda et al., Ghrelin restores ’lean-type’ hunger and energy expenditure profiles in morbidly obese subjects but has no effect on postgastrectomy subjects. Int.J Obes.(Lond) 33, 317–325 (2009).

9. M. Nakazato et al., A role for ghrelin in the central regulation of feeding. Nature 409, 194–198 (2001).

10. H. Ariyasu et al., Stomach is a major source of circulating ghrelin, and feeding state determines plasma ghrelin-like immunoreactivity levels in humans. J.Clin.Endocrinol.Metab 86, 4753–4758 (2001).

11. M. Tschop et al., Post-prandial decrease of circulating human ghrelin levels. J.Endocrinol.Invest 24, RC19–RC21 (2001).

12. I. Mandic et al., The effects of exercise and ambient temperature on dietary intake, appetite sensation, and appetite regulating hormone concentrations. Nutr Metab (Lond) 16, 29 (2019).

13. P. J. Tomasik, K. Sztefko, M. Pizon, The effect of short-term cold and hot exposure on total plasma ghrelin concentrations in humans. Horm Metab Res 37, 189–190 (2005).

14. K. Tokizawa, Y. Onoue, Y. Uchida, K. Nagashima, Ghrelin induces time-dependent modulation of thermoregulation in the cold. Chronobiol Int 29, 736–746 (2012).

15. M. S. Kim et al., Changes in ghrelin and ghrelin receptor expression according to feeding status. NeuroReport 14, 1317–1320 (2003).

16. B. Ueberberg, N. Unger, W. Saeger, K. Mann, S. Petersenn, Expression of ghrelin and its receptor in human tissues. Horm Metab Res 41, 814–821 (2009).

17. J. M. Zigman, J. E. Jones, C. E. Lee, C. B. Saper, J. K. Elmquist, Expression of ghrelin receptor mRNA in the rat and the mouse brain. J.Comp Neurol. 494, 528–548 (2006).

18. J. Kamegai et al., Central effect of ghrelin, an endogenous growth hormone secretagogue, on hypothalamic peptide gene expression. Endocrinology 141, 4797–4800 (2000).

19. M. A. Thomas, V. Ryu, T. J. Bartness, Central ghrelin increases food foraging/hoarding that is blocked by GHSR antagonism and attenuates hypothalamic paraventricular nucleus neuronal activation. Am J Physiol Regul Integr Comp Physiol, ajpregu 00216 02015 (2015).

20. M. A. Cowley et al., The distribution and mechanism of action of ghrelin in the CNS demonstrates a novel hypothalamic circuit regulating energy homeostasis. Neuron 37, 649–661 (2003).

21. J. M. Zigman et al., Mice lacking ghrelin receptors resist the development of diet-induced obesity. J.Clin.Invest 115, 3564–3572 (2005).

22. K. E. Wortley et al., Genetic deletion of ghrelin does not decrease food intake but influences metabolic fuel preference. Proc.Natl.Acad.Sci.U.S.A 101, 8227–8232 (2004).

23. J. H. Lee et al., Neuronal Deletion of Ghrelin Receptor Almost Completely Prevents Diet-Induced Obesity. Diabetes 65, 2169–2178 (2016).

24. L. Lin et al., The suppression of ghrelin signaling mitigates age-associated thermogenic impairment. Aging (Albany NY) 6, 1019–1032 (2014).

25. L. Lin et al., Ablation of ghrelin receptor reduces adiposity and improves insulin sensitivity during aging by regulating fat metabolism in white and brown adipose tissues. Aging Cell 10, 996–1010 (2011).

26. K. Dezaki et al., Blockade of pancreatic islet-derived ghrelin enhances insulin secretion to prevent high-fat diet-induced glucose intolerance. Diabetes 55, 3486–3493 (2006).

27. J. C. Chuang et al., Ghrelin mediates stress-induced food-reward behavior in mice. J Clin Invest 121, 2684–2692 (2011).

28. Q. Wang et al., Arcuate AgRP neurons mediate orexigenic and glucoregulatory actions of ghrelin. Mol.Metab 3, 64–72 (2014).

29. E. A. Davis et al., Ghrelin Signaling Affects Feeding Behavior, Metabolism, and Memory through the Vagus Nerve. Curr Biol 30, 4510–4518 e4516 (2020).

30. Y. Date et al., The role of the gastric afferent vagal nerve in ghrelin-induced feeding and growth hormone secretion in rats. Gastroenterology 123, 1120–1128 (2002).

31. C. W. Le Roux et al., Ghrelin does not stimulate food intake in patients with surgical procedures involving vagotomy. J Clin Endocrinol.Metab 90, 4521–4524 (2005).

32. A. Mano-Otagiri, H. Ohata, A. Iwasaki-Sekino, T. Nemoto, T. Shibasaki, Ghrelin suppresses noradrenaline release in the brown adipose tissue of rats. J.Endocrinol. 201, 341–349 (2009).

33. M. Arnold, A. Mura, W. Langhans, N. Geary, Gut vagal afferents are not necessary for the eating-stimulatory effect of intraperitoneally injected ghrelin in the rat. J Neurosci 26, 11052–11060 (2006).

34. A. A. Romanovsky, V. A. Kulchitsky, C. T. Simons, N. Sugimoto, M. Szekely, Cold defense mechanisms in vagotomized rats. Am J Physiol 273, R784–789 (1997).

35. S. J. Brookes, N. J. Spencer, M. Costa, V. P. Zagorodnyuk, Extrinsic primary afferent signalling in the gut. Nat Rev Gastroenterol Hepatol 10, 286–296 (2013).

36. J. Cui, J. Himms-Hagen, Rapid but transient atrophy of brown adipose tissue in capsaicin-desensitized rats. Am.J.Physiol. 262, R562–R567 (1992).

37. J. Cui, J. Himms-Hagen, Long-term decrease in body fat and in brown adipose tissue in capsaicin-desensitized rats. Am.J.Physiol. 262, R568–R573 (1992).

38. J. Cui, G. Zaror-Behrens, J. Himms-Hagen, Capsaicin desensitization induces atrophy of brown adipose tissue in rats. Am.J.Physiol. 259, R324–R332 (1990).

39. H. Fukuda et al., Ghrelin enhances gastric motility through direct stimulation of intrinsic neural pathways and capsaicin-sensitive afferent neurones in rats. Scand J Gastroenterol 39, 1209–1214 (2004).

40. H. Jiang, L. Betancourt, R. G. Smith, Ghrelin amplifies dopamine signaling by cross talk involving formation of growth hormone secretagogue receptor/dopamine receptor subtype 1 heterodimers. Mol Endocrinol 20, 1772–1785 (2006).

41. S. R. Chen et al., Ghrelin receptors mediate ghrelin-induced excitation of agouti-related protein/neuropeptide Y but not pro-opiomelanocortin neurons. J Neurochem 142, 512–520 (2017).

42. B. K. Mani et al., LEAP2 changes with body mass and food intake in humans and mice. J Clin Invest 129, 3909–3923 (2019).

43. W. Li et al., Characterization of voltage-and Ca2+-activated K+ channels in rat dorsal root ganglion neurons. J Cell Physiol 212, 348–357 (2007).

44. C. M’Kadmi et al., N-Terminal Liver-Expressed Antimicrobial Peptide 2 (LEAP2) Region Exhibits Inverse Agonist Activity toward the Ghrelin Receptor. J Med Chem 62, 965–973 (2019).

45. J. G. Verbalis, E. M. Stricker, A. G. Robinson, G. E. Hoffman, Cholecystokinin activates cFos expression in hypothalamic oxytocin and corticotropin releasing hormone neurons. Journal of Neuroendocrinology 3, 205–213 (1991).

46. S. N. Lawson, P. J. Waddell, Soma neurofilament immunoreactivity is related to cell size and fibre conduction velocity in rat primary sensory neurons. J Physiol 435, 41–63 (1991).

47. S. P. Hunt, P. W. Mantyh, The molecular dynamics of pain control. Nat Rev Neurosci 2, 83–91 (2001).

48. C. E. Le Pichon, A. T. Chesler, The functional and anatomical dissection of somatosensory subpopulations using mouse genetics. Front Neuroanat 8, 21 (2014).

49. X. Zhou et al., Deletion of PIK3C3/Vps34 in sensory neurons causes rapid neurodegeneration by disrupting the endosomal but not the autophagic pathway. Proc Natl Acad Sci U S A 107, 9424–9429 (2010).

50. M. M. Scott, et al., Hindbrain ghrelin receptor signaling is sufficient to maintain fasting glucose. PLoS.ONE. 7, e44089 (2012).

51. Y. Sun, N. F. Butte, J. M. Garcia, R. G. Smith, Characterization of adult ghrelin and ghrelin receptor knockout mice under positive and negative energy balance. Endocrinology 149, 843–850 (2008).

52. F. Naznin et al., Diet-induced obesity causes peripheral and central ghrelin resistance by promoting inflammation. J Endocrinol 226, 81–92 (2015).

53. A. I. Mina et al., CalR: A Web-Based Analysis Tool for Indirect Calorimetry Experiments. Cell Metab 28, 656–666 e651 (2018).

54. B. Cannon, J. Nedergaard, Nonshivering thermogenesis and its adequate measurement in metabolic studies. J.Exp.Biol. 214, 242–253 (2011).

55. H. M. Feldmann, V. Golozoubova, B. Cannon, J. Nedergaard, UCP1 ablation induces obesity and abolishes diet-induced thermogenesis in mice exempt from thermal stress by living at thermoneutrality. Cell Metab 9, 203–209 (2009).

56. B. N. Gaskill et al., Heat or insulation: behavioral titration of mouse preference for warmth or access to a nest. PLoS One 7, e32799 (2012).

57. T. G. Youngstrom, T. J. Bartness, White adipose tissue sympathetic nervous system denervation increases fat pad mass and fat cell number. Am.J.Physiol. 275, R1488–R1493 (1998).

58. R. D. Dix, R. R. McKendall, J. R. Baringer, Comparative neurovirulence of herpes simplex virus type 1 strains after peripheral or intracerebral inoculation of BALB/c mice. Infect.Immun. 40, 103–112 (1983).

59. G. L. Demmin, A. C. Clase, J. A. Randall, L. W. Enquist, B. W. Banfield, Insertions in the gG gene of pseudorabies virus reduce expression of the upstream Us3 protein and inhibit cell-to-cell spread of virus infection. J.Virol. 75, 10856–10869 (2001).

60. B. N. Smith et al., Pseudorabies virus expressing enhanced green fluorescent protein: A tool for in vitro electrophysiological analysis of transsynaptically labeled neurons in identified central nervous system circuits. Proc.Natl.Acad.Sci U.S.A 97, 9264–9269 (2000).

61. N. L. Nguyen et al., Separate and shared sympathetic outflow to white and brown fat coordinately regulates thermoregulation and beige adipocyte recruitment. Am J Physiol Regul Integr Comp Physiol 312, R132–R145 (2017).

62. V. Ryu, A. G. Watts, B. Xue, T. J. Bartness, Bidirectional crosstalk between the sensory and sympathetic motor systems innervating brown and white adipose tissue in male Siberian hamsters. Am J Physiol Regul Integr Comp Physiol 312, R324–R337 (2017).

63. T. D. Muller et al., Ghrelin. Mol Metab 4, 437–460 (2015).

64. P. T. Pfluger et al., Simultaneous deletion of ghrelin and its receptor increases motor activity and energy expenditure. Am.J.Physiol Gastrointest.Liver Physiol 294, G610–G618 (2008).

65. K. N. Browning, R. A. Travagli, Central nervous system control of gastrointestinal motility and secretion and modulation of gastrointestinal functions. Compr Physiol 4, 1339–1368 (2014).

66. V. Ryu, T. J. Bartness, Short and long sympathetic-sensory feedback loops in white fat. Am.J.Physiol Regul.Integr.Comp Physiol, (2014).

67. V. Ryu, J. T. Garretson, Y. Liu, C. H. Vaughan, T. J. Bartness, Brown adipose tissue has sympathetic-sensory feedback circuits. J Neurosci 35, 2181–2190 (2015).

68. E. Ebert, Gastrointestinal involvement in spinal cord injury: a clinical perspective. J Gastrointestin Liver Dis 21, 75–82 (2012).

69. M. Bamshad, V. T. Aoki, M. G. Adkison, W. S. Warren, T. J. Bartness, Central nervous system origins of the sympathetic nervous system outflow to white adipose tissue. Am.J.Physiol. 275, R291–R299 (1998).

70. T. J. Bartness, Y. B. Shrestha, C. H. Vaughan, G. J. Schwartz, C. K. Song, Sensory and sympathetic nervous system control of white adipose tissue lipolysis. Mol.Cell Endocrinol. 318, 34–43 (2010).

71. H. Munzberg, E. Qualls-Creekmore, H. R. Berthoud, C. D. Morrison, S. Yu, Neural Control of Energy Expenditure. Handb Exp Pharmacol 233, 173–194 (2016).

72. K. Rezai-Zadeh, H. Munzberg, Integration of sensory information via central thermoregulatory leptin targets. Physiol Behav 121, 49–55 (2013).

73. H. M. Young, D. Ciampoli, J. Hsuan, A. J. Canty, Expression of Ret-, p75(NTR)-, Phox2a-, Phox2b-, and tyrosine hydroxylase-immunoreactivity by undifferentiated neural crest-derived cells and different classes of enteric neurons in the embryonic mouse gut. Dev Dyn 216, 137–152 (1999).

74. A. Pattyn, X. Morin, H. Cremer, C. Goridis, J. F. Brunet, The homeobox gene Phox2b is essential for the development of autonomic neural crest derivatives. Nature 399, 366–370 (1999).

75. G. Paxinos, K. B. J. Franklin, The Mouse Brain in Stereotaxic Coordinates. (Academic Press, New York, ed. 2nd, 2007).

76. J. T. Garretson et al., Lipolysis sensation by white fat afferent nerves triggers brown fat thermogenesis. Mol Metab 5, 626–634 (2016).

77. G. Paxinos, K. B. J. Franklin, The mouse brain in stereotaxic coordinates. (Academic Press, San Diego, 2001).

78. E. Frey et al., An in vitro assay to study induction of the regenerative state in sensory neurons. Exp Neurol 263, 350–363 (2015).

79. S. Estes, L. R. Zhong, L. Artinian, K. Tornieri, V. Rehder, The role of action potentials in determining neuron-type-specific responses to nitric oxide. Dev Neurobiol 75, 435–451 (2015).

80. B. F. Grewe, D. Langer, H. Kasper, B. M. Kampa, F. Helmchen, High-speed in vivo calcium imaging reveals neuronal network activity with near-millisecond precision. Nat Methods 7, 399–405 (2010).

81. G. Grynkiewicz, M. Poenie, R. Y. Tsien, A new generation of Ca2+ indicators with greatly improved fluorescence properties. J Biol Chem 260, 3440–3450 (1985).

82. R. W. Teichert et al., Functional profiling of neurons through cellular neuropharmacology. Proc Natl Acad Sci U S A 109, 1388–1395 (2012).

83. S. A. Thayer, R. J. Miller, Regulation of the intracellular free calcium concentration in single rat dorsal root ganglion neurones in vitro. J Physiol 425, 85–115 (1990).

84. L. R. Zhong, S. Estes, L. Artinian, V. Rehder, Acetylcholine elongates neuronal growth cone filopodia via activation of nicotinic acetylcholine receptors. Dev Neurobiol 73, 487–501 (2013).

85. L. Artinian, K. Tornieri, L. Zhong, D. Baro, V. Rehder, Nitric oxide acts as a volume transmitter to modulate electrical properties of spontaneously firing neurons via apamin-sensitive potassium channels. J Neurosci 30, 1699–1711 (2010).

86. L. R. Zhong, L. Artinian, V. Rehder, Dopamine suppresses neuronal activity of Helisoma B5 neurons via a D2-like receptor, activating PLC and K channels. Neuroscience 228, 109–119 (2013).

87. L. R. Zhong, S. Estes, L. Artinian, V. Rehder, Nitric oxide regulates neuronal activity via calcium-activated potassium channels. PLoS One 8, e78727 (2013).

88. K. J. Livak, T. D. Schmittgen, Analysis of relative gene expression data using real-time quantitative PCR and the 2(-Delta Delta C(T)) Method. Methods 25, 402–408 (2001).

89. Y. Cui, T. F. Lee, L. C. H. Wang, In vivo microdialysis study on changes in septal dynorphin and á- endorphin activities in active and hibernating Columbian ground squirrels. Brain Research 710, 271–274 (1996).

90. C. H. Vaughan, E. Zarebidaki, J. C. Ehlen, T. J. Bartness, Analysis and measurement of the sympathetic and sensory innervation of white and brown adipose tissue. Methods Enzymol. 537, 199–225 (2014).

91. B. K. Mani, S. Osborne-Lawrence, P. Vijayaraghavan, C. Hepler, J. M. Zigman, beta1-Adrenergic receptor deficiency in ghrelin-expressing cells causes hypoglycemia in susceptible individuals. J Clin Invest 126, 3467–3478 (2016).

