## Supplemental Figures, Figure Legends and Tables for "Crosstalk between Gut Sensory Ghrelin Signaling and Adipose Tissue Sympathetic Outflow Regulates Metabolic Homeostasis"

Department of Biology, Georgia State University, 24 Peachtree Center Avenue, Atlanta, GA 30303, USA

Department of Biology, Georgia State University, 24 Peachtree Center Avenue, Atlanta, GA 30303, USA

Running Title: Peripheral sensory ghrelin signaling regulates energy homeostasis.

#### Supplemental Figures and Figure Legends

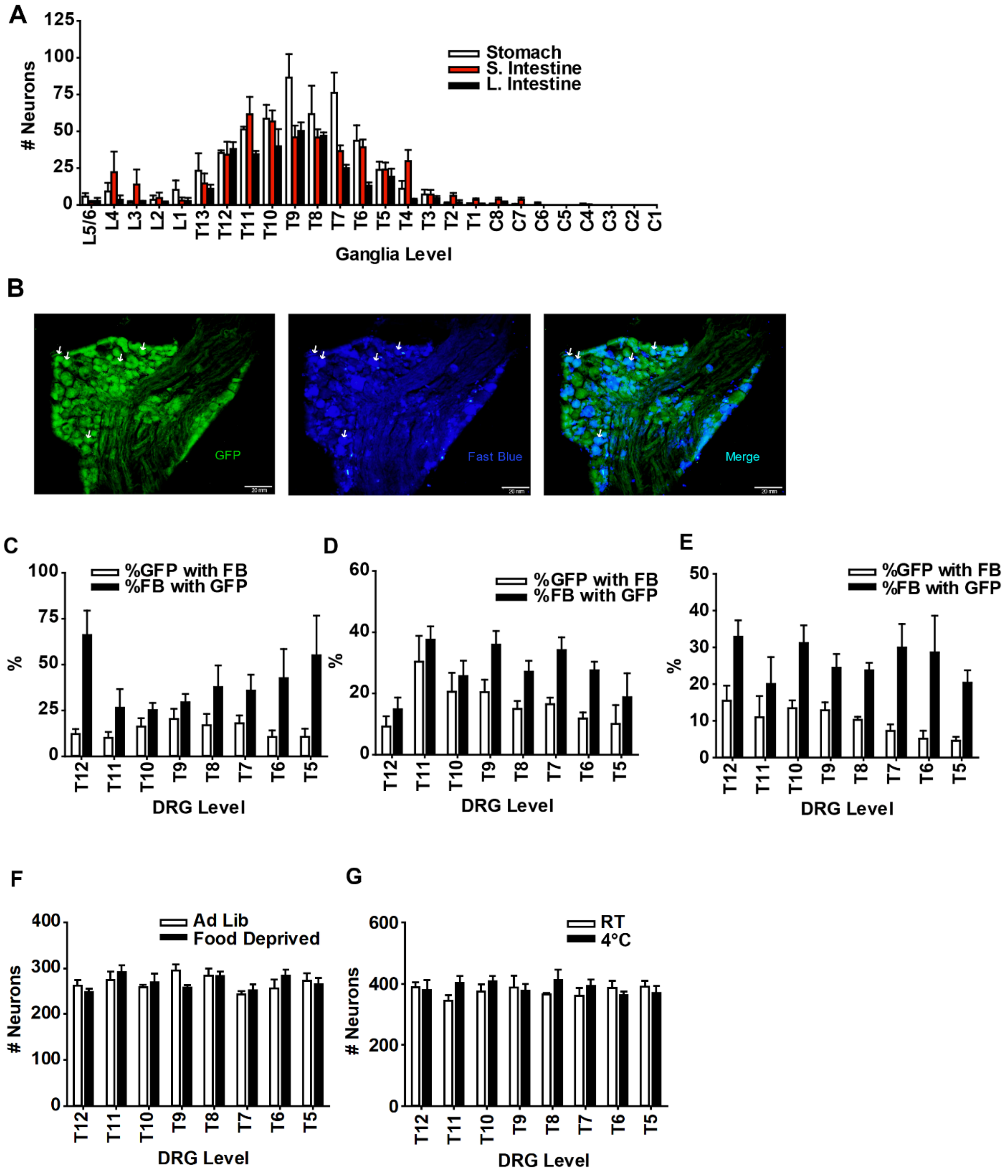

**Supplemental Figure 1. DRG neurons express GHSR and project to the gastrointestinal system.**

(A) Number of Fast Blue-positive neurons following injection into the stomach, small intestine, or large intestine in L5/6-C1 level DRGs; n = 8 mice/group.

(B) Representative GHSR-GFP, Fast Blue and GFP/Fast Blue double labeling in T10 level DRG from GHSR-tau-GFP mice. Fast Blue was injected into the stomach.

(C) % of GHSR-GFP- and Fast Blue-positive neurons in T5-T12 level DRGs from GHSR-tau-GFP mice. Fast Blue was injected into the stomach. n = 8 mice/group.

(D) % of GHSR-GFP- and Fast Blue-positive neurons in T5-T12 level DRGs from GHSR-tau-GFP mice. Fast Blue was injected into the small intestine. n = 8 mice/group.

(E) % of GHSR-GFP- and Fast Blue-positive neurons in T5-T12 level DRGs from GHSR-tau-GFP mice. Fast Blue was injected into the large intestine. n = 8 mice/group.

(F) Number of GHSR-GFP neurons in *ad lib*-fed and overnight food deprived mice; n = 8 mice/group.

(G) Number of GHSR-GFP neurons in room temperature and 1-day 4°C cold exposed mice; n = 8 mice/group.

Data are presented as mean ± SEM. * $p<0.05$ as measured by two-tailed unpaired t-test.

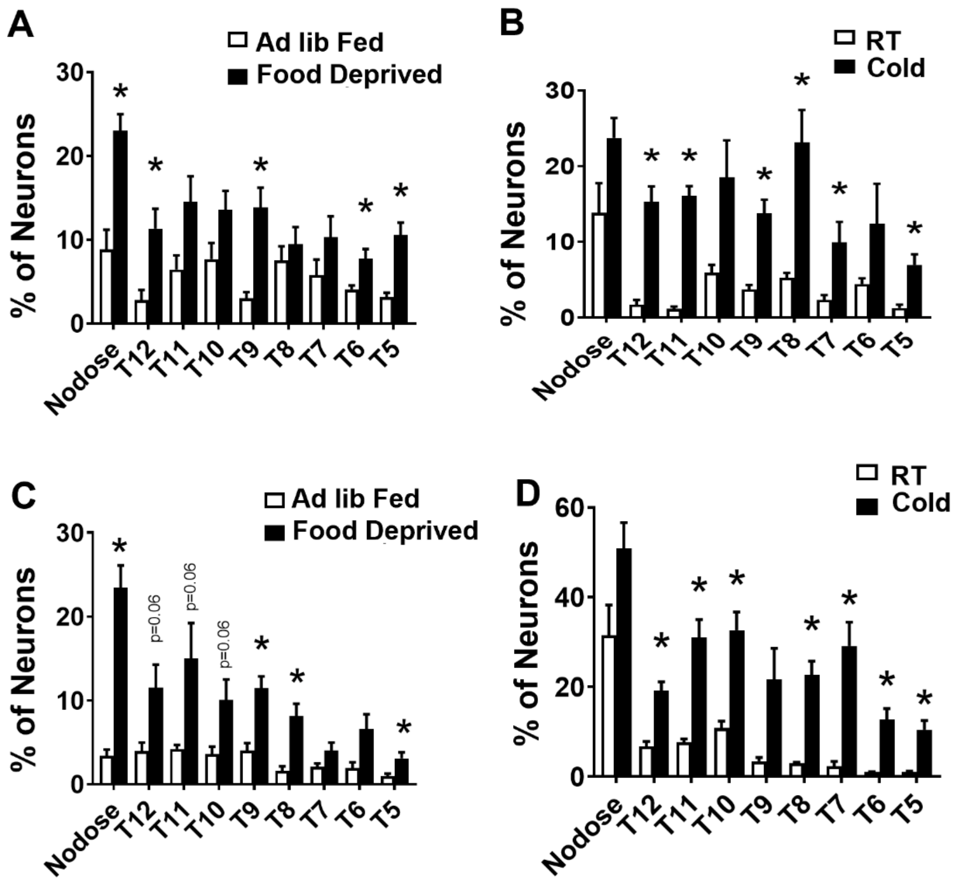


**Supplemental Figure 2. DRG GHSR-expressing neurons are regulated by energetic challenges.**

(A) Percentage of triple-labeled (GFP, cFos, and Fast Blue) neurons from nodose and T12-T5 level DRGs in stomach Fast Blue-injected mice following overnight food deprivation; n = 4 mice/group.

(B) Percentage of triple-labeled (GFP, cFos, and Fast Blue) neurons from nodose and T12-T5 level DRGs in stomach Fast Blue-injected mice following 1-day 4°C cold exposure; n = 4 mice/group.

(C) Percentage of triple-labeled (GFP, cFos, and Fast Blue) neurons from nodose and T12-T5 level DRGs in small intestine Fast Blue-injected mice following overnight food deprivation; n = 4 mice/group.

(D) Percentage of triple-labeled (GFP, cFos, and Fast Blue) neurons from nodose and T12-T5 level DRGs in small intestine Fast Blue-injected mice following 1-day 4°C cold exposure; n = 4 mice/group.

Data are presented as mean ± SEM. *, $p<0.05$ as measured by two-tailed unpaired t-test.

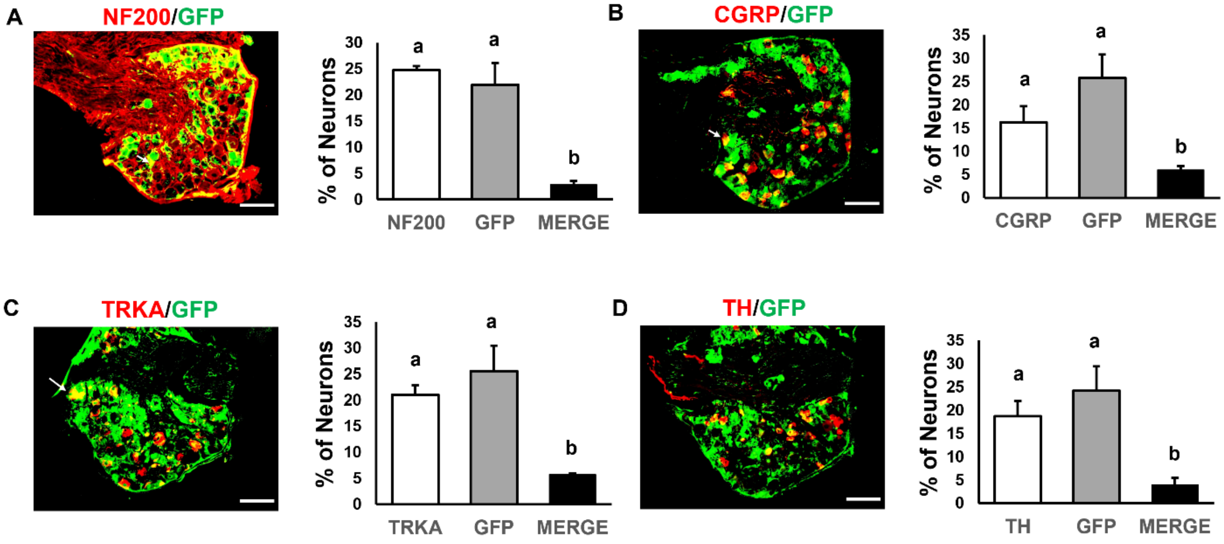


**Supplemental Figure 3. Characterization of DRG GHSR-positive neurons.**

(A) Representative image and percentage of NF200, GFP, and double-labeled neurons in DRG of 2-month-old GHSR-Tau-GFP mice. n = 3 mice/group.
(B) Representative image and percentage of CGRP, GFP, and double-labeled neurons in DRG of 2-month-old GHSR-Tau-GFP mice. n = 3 mice/group.
(C) Representative image and percentage of TRKA, GFP, and double-labeled neurons in DRG of 2-month-old GHSR-Tau-GFP mice. n = 3 mice/group.
(D) Representative image and percentage of TH, GFP, and double-labeled neurons in DRG of 2-month-old GHSR-Tau-GFP mice. n = 3 mice/group.

Data are presented as mean ± SEM. Bars with different lowercase letters are statistically different from each other. Significance was measured by one-way ANOVA with Bonferonni post-hoc analysis.

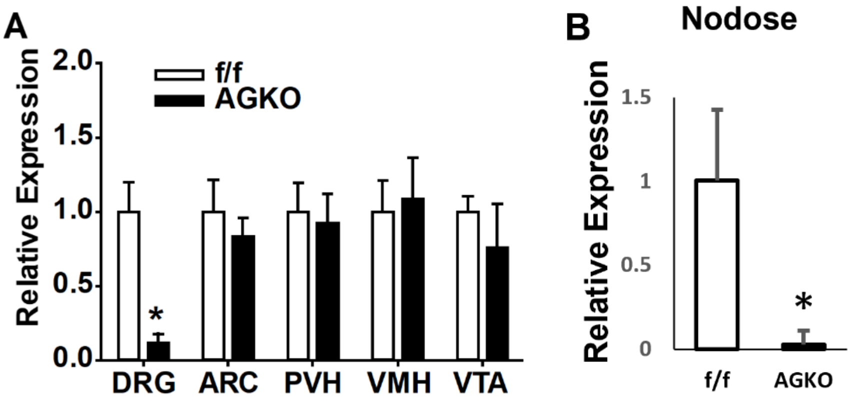


**Supplemental Figure 4. GHSR expression in AGKO and f/f mice.**

(A-B) Relative GHSR expression in DRGs, hypothalamic nuclei (arcuate hypothalamus(ARC), paraventricular hypothalamus (PVH), ventromedial hypothalamus (VMH)), and the ventral tegmental area (VTA) (A) and in the nodose ganglia (B) in f/f and AGKO mice; n = 3-6 mice/group.

Data are presented as mean ± SEM. *, $p<0.05$ as measured by two-tailed unpaired t-test.

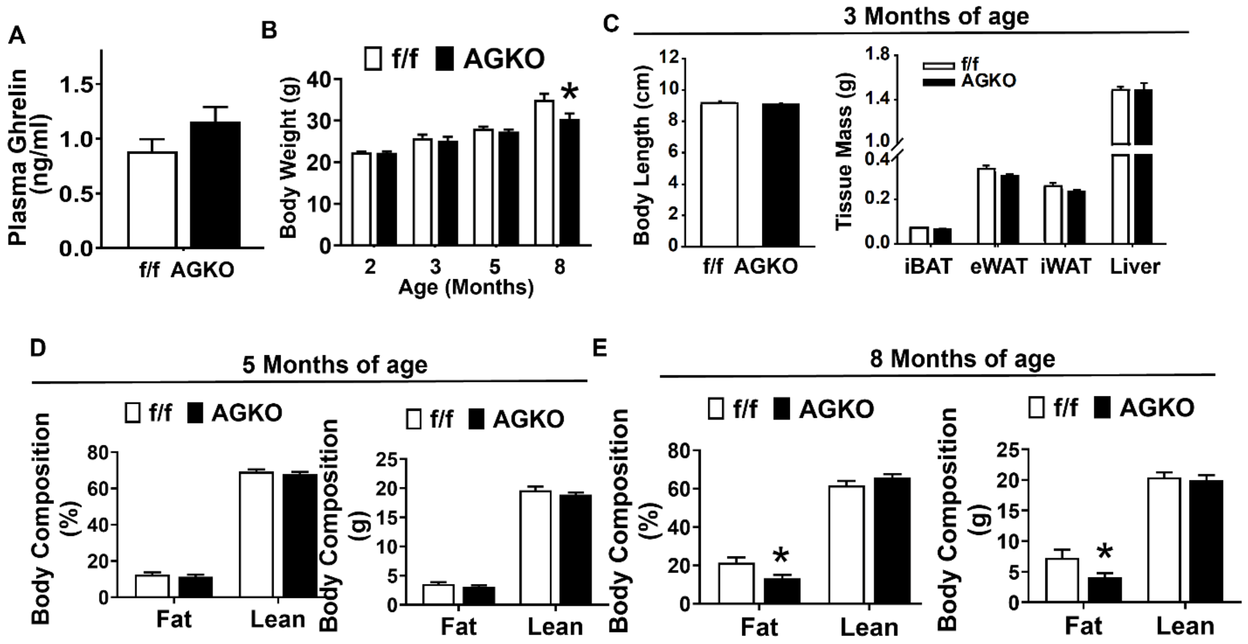


**Supplemental Figure 5. Body weight and body composition in chow-fed f/f and AGKO mice.**

(A) Plasma acyl-ghrelin levels in 2-month-old f/f and AGKO mice fed a regular chow diet. n = 4 mice/group.

(B) Body weights in chow-fed f/f and AGKO mice measured at 2 months, 3 months, 5 months and 8 months of age. n = 4 mice/group for 2 months and 3 months of age, 12 (f/f) and 9 (AGKO) for 5 months of age, and 3 (f/f) and 4 (AGKO) for 8 months of age.

(C) Body length and tissue mass (iBAT, eWAT, iWAT and liver) in chow-fed f/f and AGKO mice at 3 months of age. n = 7-8 mice/group.

(D) Body composition (% and total fat and lean mass) in chow-fed f/f and AGKO mice at 5 months of age. n = 8 mice/group.

(E) Body composition (% and total fat and lean mass) in chow-fed f/f and AGKO mice at 5 months of age. n = 4 mice/group.

Data are presented as mean ± SEM. *, $p<0.05$ as measured by two-tailed unpaired t-test.

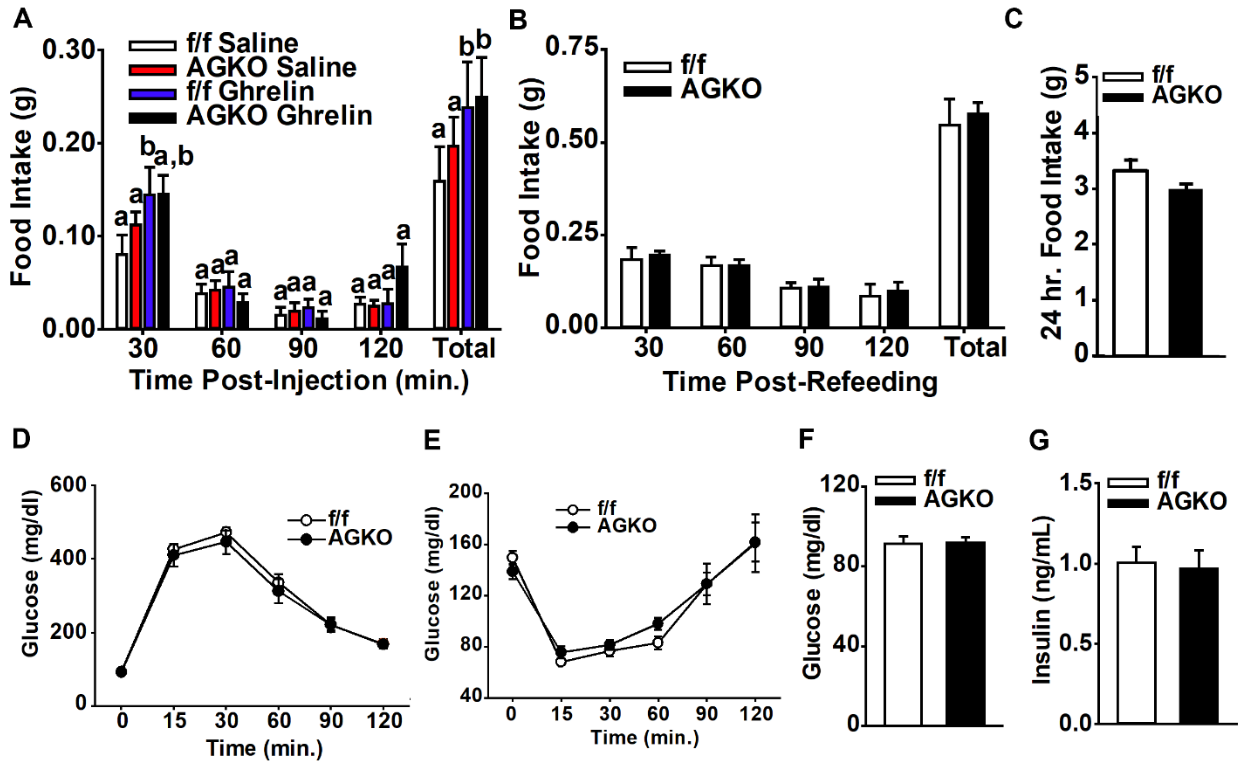


**Supplemental Figure 6. Metabolic phenotyping in chow-fed f/f and AGKO mice.**

(A) Food intake measurements following subcutaneous saline or ghrelin (0.5mg/kg body mass) injections in chow-fed f/f and AGKO mice; n = 8 mice/group.
(B) Food intake measurements following overnight fasting in chow-fed f/f and AGKO mice; n = 8 mice/group.
(C) Twenty-four-hour *ad lib* food intake in chow-fed f/f and AGKO mice; n = 8 mice/group.
(D) Glucose tolerance test in chow fed f/f and AGKO mice; n = 7-8 mice/group.
(E) Insulin tolerance test in chow fed f/f and AGKO mice; n = 7-8 mice/group.
(F) Fasting blood glucose levels in chow fed f/f and AGKO mice; n = 7-8 mice/group.
(G) Blood insulin levels in *ad lib* chow fed f/f and AGKO mice; n = 7-8 mice/group.

Data are presented as mean ± SEM. In (A), bars with the same lowercase letters are not statistically different, and bars with different lowercase letters signify statistical significance at $p<0.05$. Statistical significance was measured by unpaired two-tailed t-test or two-way ANOVA with Bonferonni post-hoc analysis.

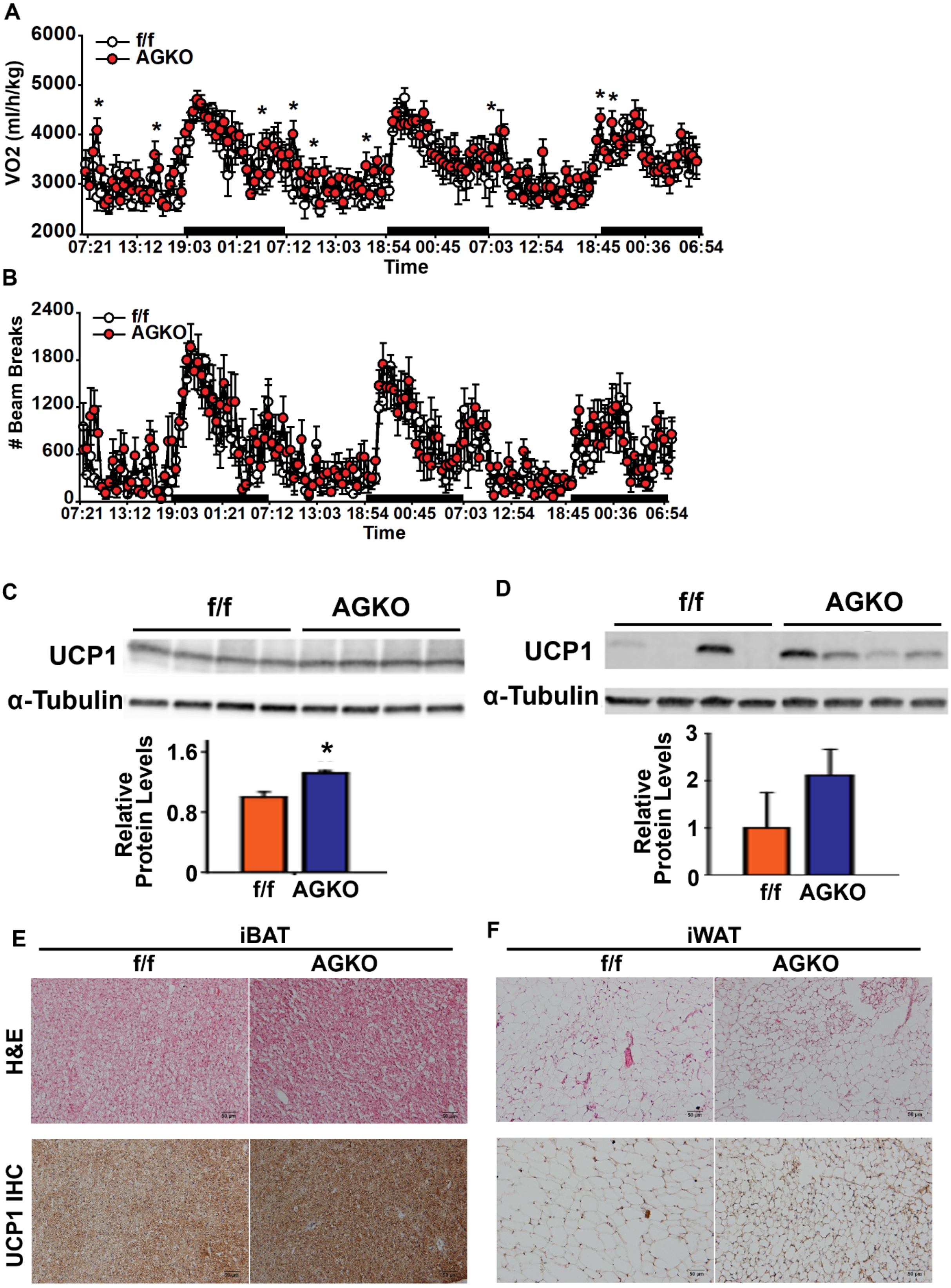
A
VO2 (ml/h/kg)
f/f
AGKO
Time
B
### Beam Breaks
f/f
AGKO
Time
C
f/f
AGKO
UCP1
α-Tubulin
Relative Protein Levels
f/f
AGKO
D
f/f
AGKO
UCP1
α-Tubulin
Relative Protein Levels
f/f
AGKO
E
iBAT
f/f
AGKO
H&E
UCP1 IHC
F
iWAT
f/f
AGKO

**Supplemental Figure 7. Chow-fed sensory neuron GHSR knockout mice have slightly increased energy expenditure.**

(A) Average oxygen consumption rate in daytime and nighttime in 12-week chow-fed f/f and AGKO mice; n = 7 mice/group.

(B) Average locomotor activity during light (day) and dark (night) cycles in 12-week chow-fed f/f and AGKO mice; n = 7 mice/group.

(C)-(D) UCP1 protein levels in iBAT (C) and iWAT (D) in 12-week chow-fed f/f and AGKO mice; n = 4 mice/group.

(E)-(F) Representative H&E staining and UCP1 IHC staining in iBAT (E) and iWAT (F) of 12-week chow-fed f/f and AGKO mice.

Data are presented as mean ± SEM. * $p<0.05$ versus f/f controls as measured by two-tailed unpaired t-test.

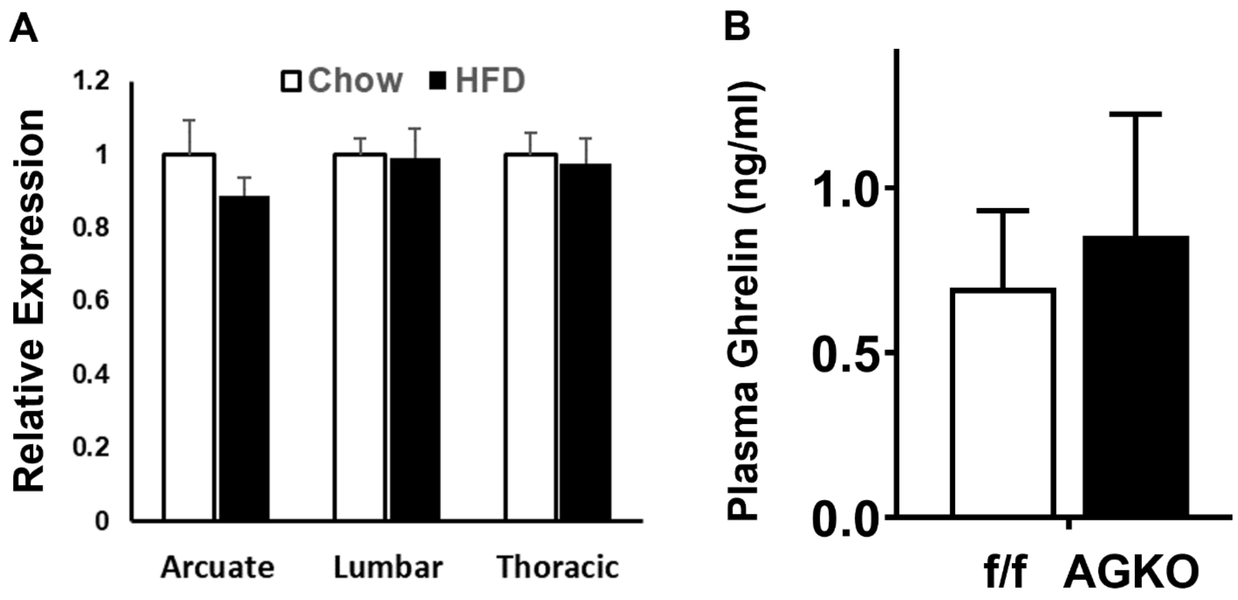


**Supplemental Figure 8. GHSR expression in arcuate hypothalamus and lumbar and thoracic DRGs in animals fed with chow or HFD for 4 weeks (A) and plasma acyl-ghrelin levels in f/f and AGKO mice fed HFD (B).** n = 7-8 mice/group in (A) and n = 3 mice/group in (B).

Data are presented as mean ± SEM.

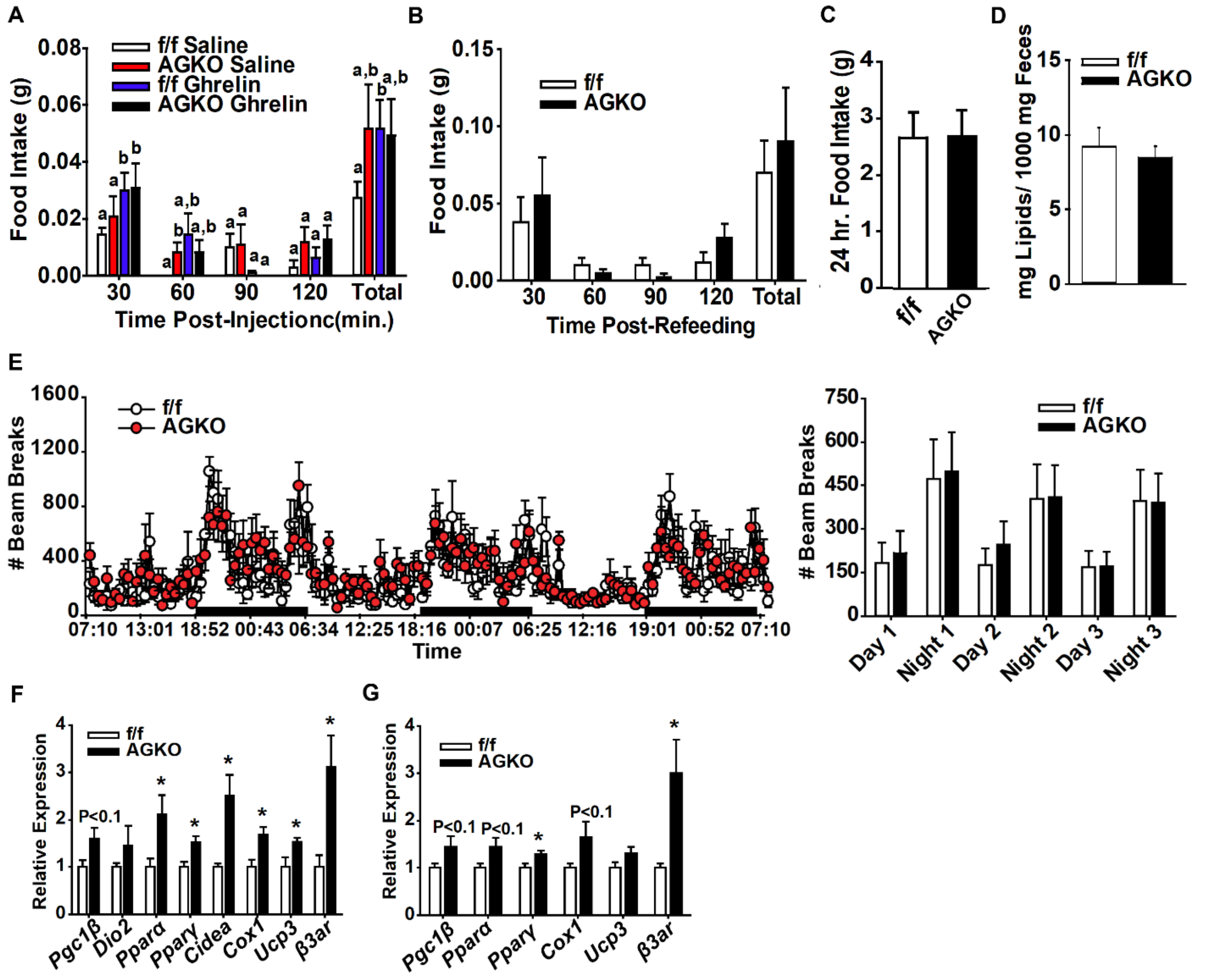


**Supplemental Figure 9. Metabolic phenotyping in f/f and AGKO mice fed HFD.**

(A) Food intake measurements following subcutaneous saline or ghrelin (0.5mg/kg body mass) injections in f/f and AGKO mice fed HFD for 10 weeks; n = 8 mice/group.
(B) Food intake measurements following overnight fasting in f/f and AGKO mice fed HFD for 12 weeks; n = 8 mice/group.
(C) Twenty-four-hour *ad lib* food intake in f/f and AGKO mice fed HFD for 11 weeks; n = 8 mice/group.
(D) Twenty-four-hour fecal lipid concentrations in f/f and AGKO mice fed HFD for 10 weeks; n = 8 mice/group.
(E) Average locomotor activity during daytime and nighttime in f/f and AGKO mice fed HFD for 16 weeks; n = 7 mice/group.
(F)-(G) Relative expression of thermogenic genes in eWAT (F) and iWAT (G) of f/f and AGKO mice fed 60% HFD for 17 weeks; n = 8 mice/group.

Data are presented as mean ± SEM. In (A), bars with the same lowercase letters are not statistically different, and bars with different lowercase letters signify statistical significance

($p<0.05$). * $p<0.05$ versus f/f controls. Significance was measured by two-tailed unpaired t-test or by two-way ANOVA with Bonferonni post-hoc analysis.

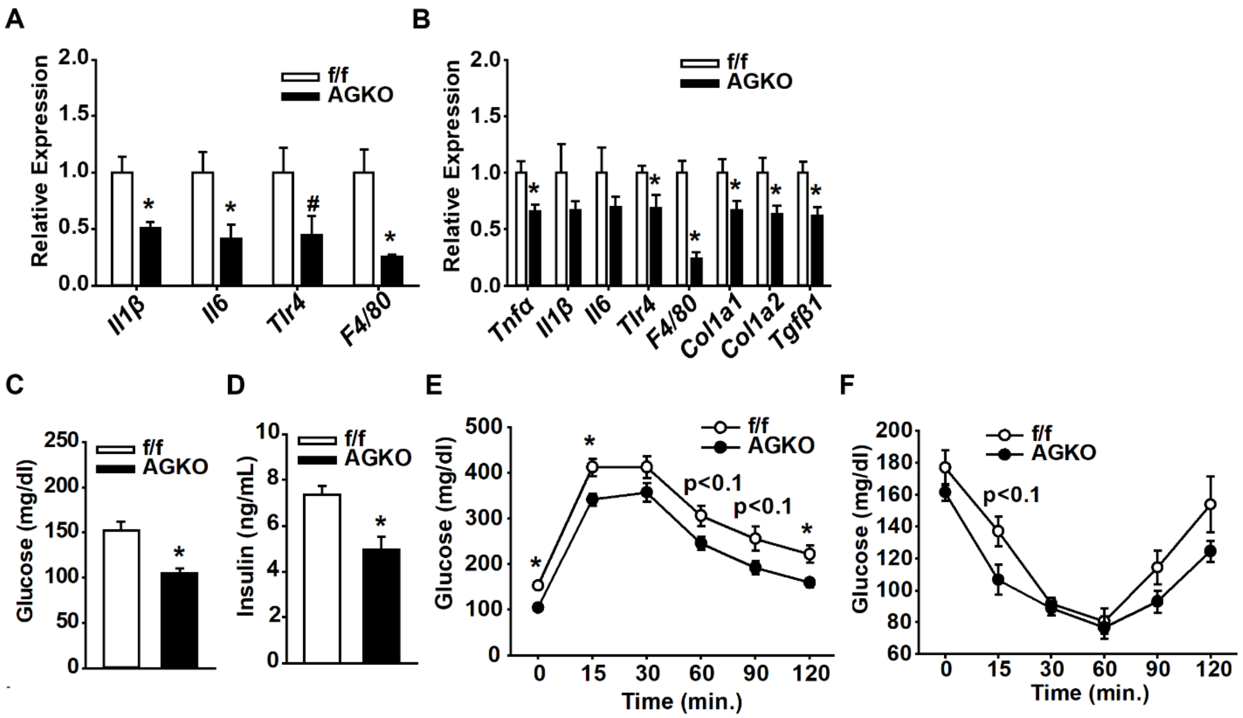


**Supplemental Figure 10. HFD-fed sensory neuron GHSR knockout mice have decreased adipose tissue inflammatory gene expression and improved insulin sensitivity.**

(A)-(B) Relative expression of inflammatory genes in iBAT (A) and eWAT (B) of f/f and AGKO mice fed HFD for 17 weeks; n = 8 mice/group.

(C) Fasting glucose levels in f/f and AGKO mice fed HFD for 14 weeks; n = 8 mice/group.

(D) Blood insulin levels in *ad lib* f/f and AGKO mice fed HFD for 15 weeks; n = 8 mice/group.

(E) Glucose tolerance test in f/f and AGKO mice fed HFD for 14 weeks; n = 8 mice/group.

(F) Insulin tolerance test in f/f and AGKO mice fed HFD for 15 weeks; n = 8 mice/group.

Data are presented as mean ± SEM. *, $p<0.05$ versus f/f controls as measured by two-tailed unpaired t-test or by two-way ANOVA with Bonferonni post-hoc analysis.

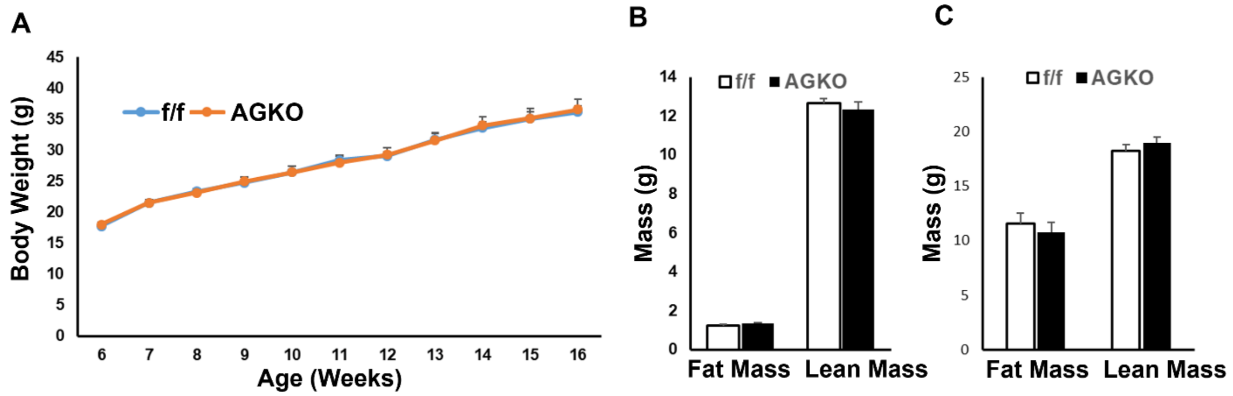


**Supplemental Figure 11. There is no difference in body weight and body composition in HFD-fed f/f and AGKO mice housed at thermoneutrality.**

(A) Weekly body weight measurements in f/f and AGKO mice fed a 60% HFD under thermoneutrality. n = 9 mice/group.

(B) –(C) Body composition in f/f and AGKO mice at 6 weeks of age before the 60% HFD feeding (B) and after fed a 60% HFD under thermoneutrality for 10 weeks (C). n = 9 mice/group.

Data are presented as mean ± SEM.

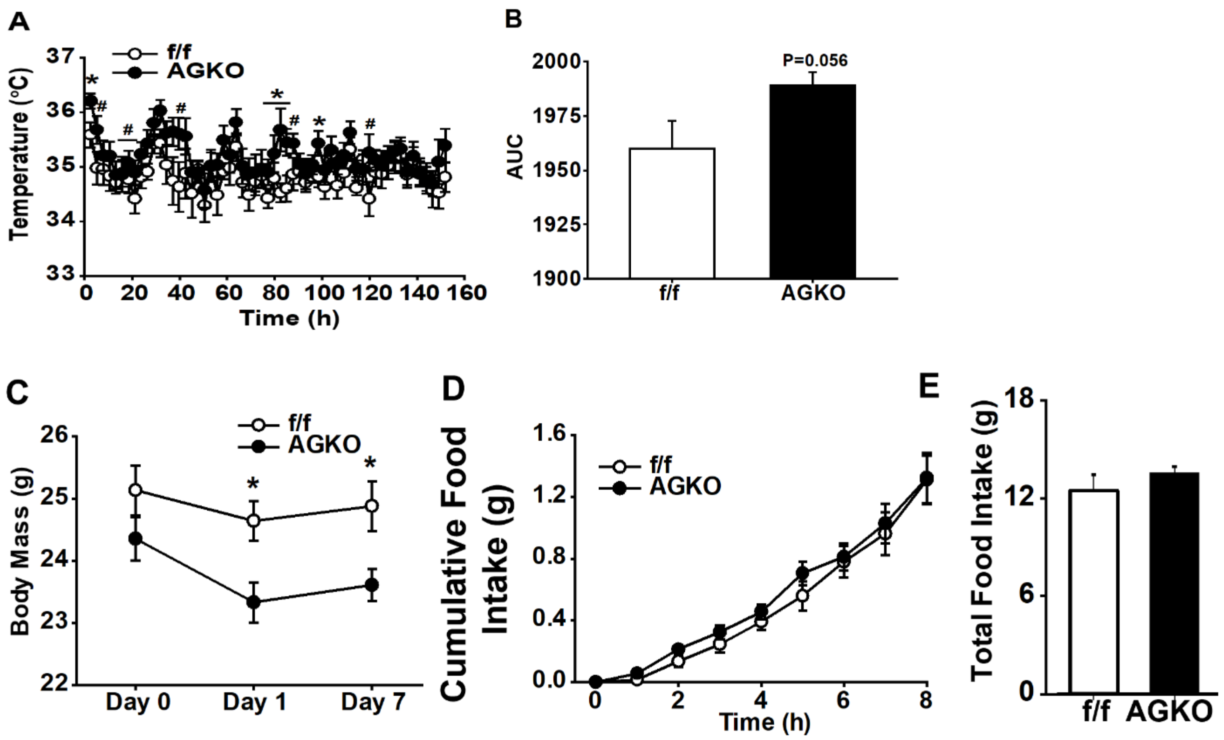


**Supplemental Figure 12. Metabolic phenotyping in f/f and AGKO mice before and after cold exposure.**

(A)-(B) Body temperature measurements (A) and area under the curve (B) from 7-day room temperature housed f/f and AGKO mice; n = 10 mice/group.

(C) Body mass measurements before cold exposure (Day 0), after 1-day cold exposure (Day 1) and 7-day cold exposure (Day 7) in f/f and AGKO mice; n = 5-7 mice/group.

(D) Cumulative food intake during the first 8-hour cold exposure in f/f and AGKO mice; n = 8 mice/group.

(E) Total food intake across 72 hours cold exposure in f/f and AGKO mice; n = 8 mice/group.

Data are presented as mean ± SEM. *, $p<0.05$ versus f/f controls as measured by two-tailed unpaired t-test or by two-way ANOVA with Bonferonni post-hoc analysis.

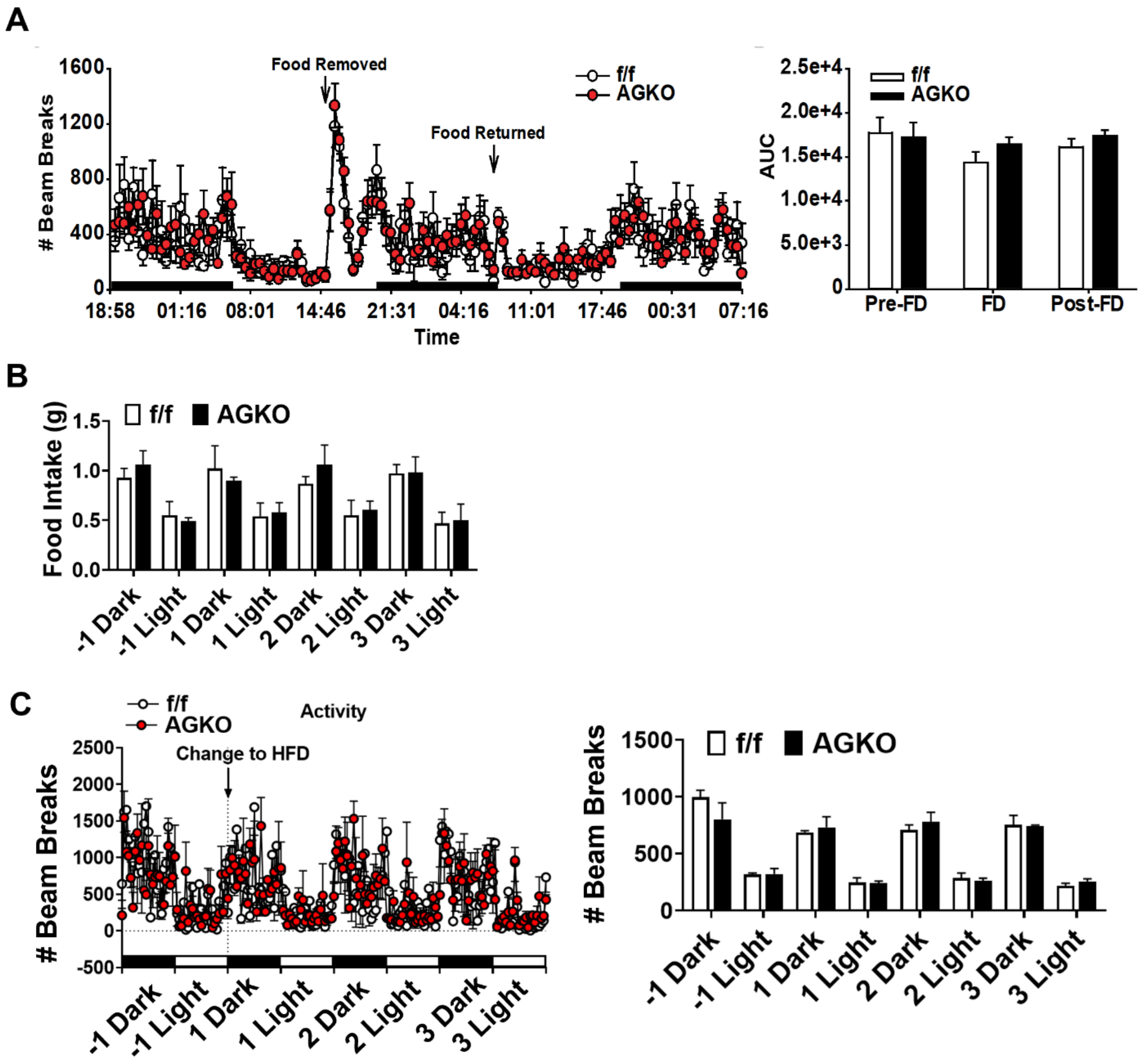


**Supplemental Figure 13. Metabolic phenotypes of sensory neuron GHSR knockout mice in response to fasting/refeeding challenge and during diet switch from chow to HFD.**

(A) Locomotor activity before, during, and after an overnight food deprivation challenge (Pre-FD, FD and Post-FD, respectively) in f/f and AGKO mice fed HDF for 12 weeks; n = 7 mice/group.

(B) Food intake during dark and light cycle 1 day before (-1 Dark and -1 Light) or 1, 2, 3 days after diet switch from LFD to HFD in 12-week f/f and AGKO mice; n = 4 mice/group.

(C) Locomotor activity during dark and light cycle 1 day before (-1 Dark and -1 Light) or 1, 2 3 days after diet switch from LFD to HFD in 12-week f/f and AGKO mice; n = 4 mice/group.

Data are presented as mean ± SEM.

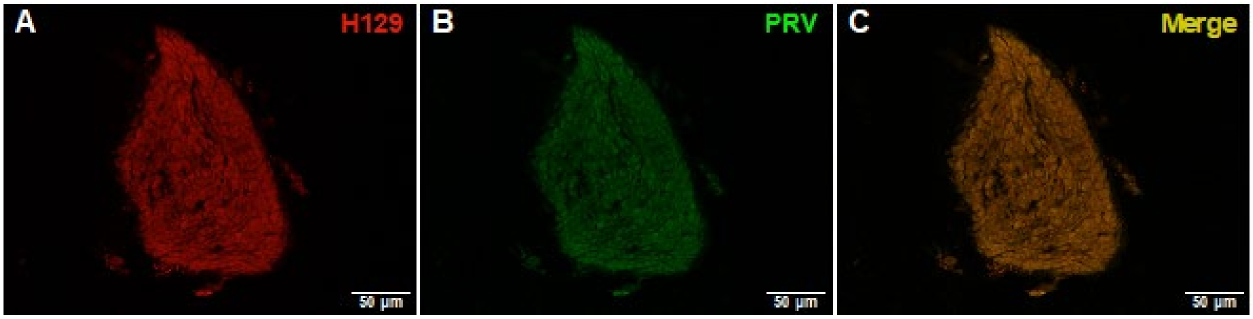


**Supplemental Figure 14. Vagotomy blocks stomach-injected H129 and iBAT-injected PRV labeling in nodose ganglia.**
Representative H129 (A), PRV (B) and double (C) labeling in the nodose ganglia following subdiaphragmatic vagotomy.

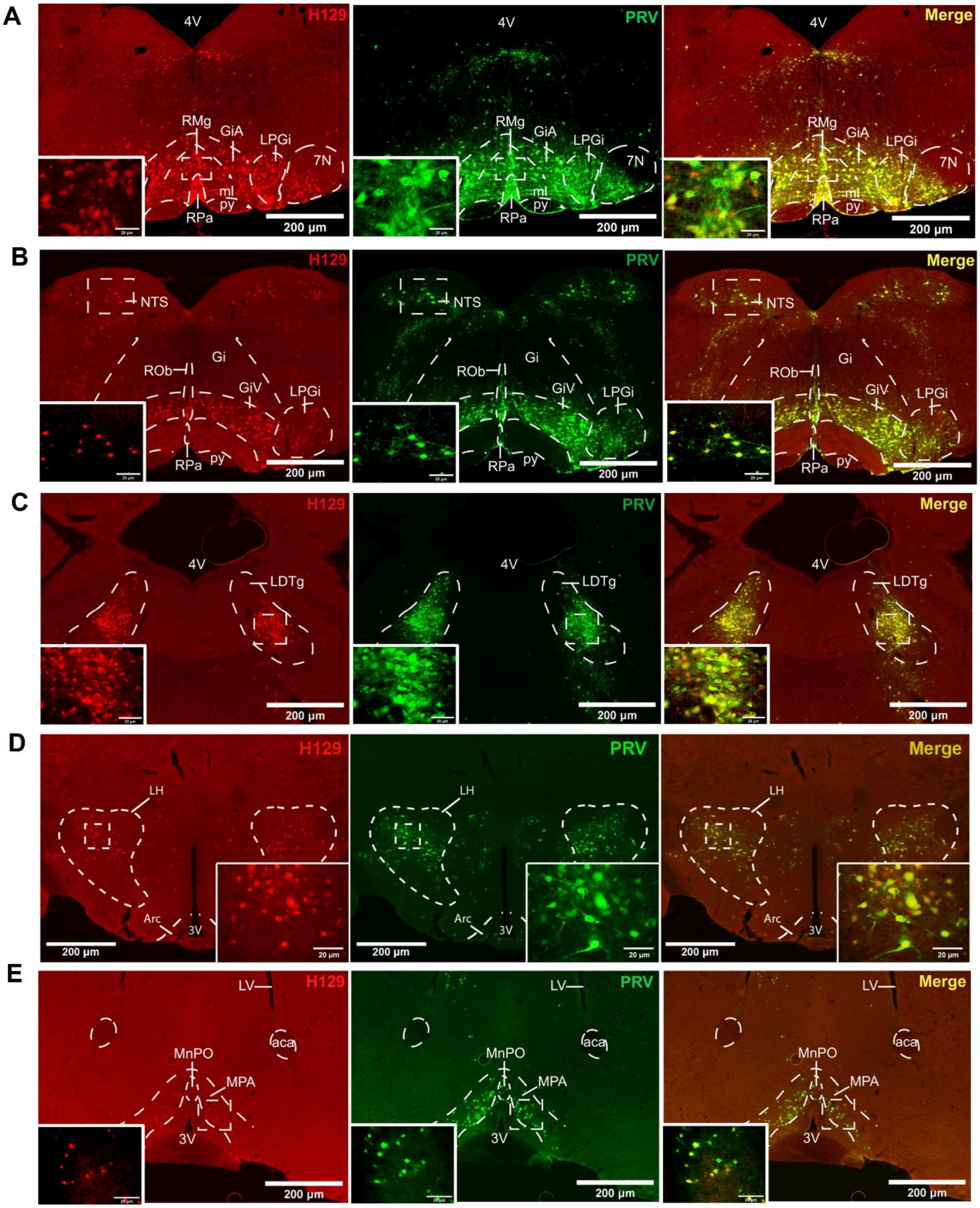
A
B
C
D
E
H129
PRV
Merge
4V
RMg
GiA
LPGi
7N
ml
py
RPa
NTS
ROb
Gi
GiV
LDTg
LH
Arc
3V
LV
aca
MnPO
MPA
200 μm
20 μm

**Supplemental Figure 15. Neuronal crosstalk between stomach-projecting DRG sensory neurons and iBAT-projecting sympathetic outflow neurons.**

(A) Representative images of H129, PRV and double labeling in hindbrain (dashed line denotes raphe magnus nucleus (RMg), raphe pallidus nucleus (RPa), gigantocellular reticular nucleus, alpha part (GiA), medial lemniscus (ml), pyramidal nucleus (py), lateral paragigantocellular nucleus (LPGi) and facial nucleus (7N); dashed box denotes area in 20x inset). 4V: $4^{th}$ ventricle.

(B) Representative images of H129, PRV and double labeling in hindbrain (dashed line denotes nucleus of the solitary tract (NTS), raphe obscurus nucleus (ROb), raphe pallidus nucleus (RPa), gigantocellular reticular nucleus, ventral part (GiV), pyramidal nucleus (py) and lateral paragigantocellular nucleus (LPGi); dashed box denotes area in 20x inset).

(C) Representative image of H129, PRV and double labeling in midbrain (dashed line denotes laterodorsal tegmental nucleus (LDTg); dashed box denotes area in 20x inset). 4V: 4th ventricle.

(D) Representative image of H129, PRV and double labeling in forebrain (dashed line denotes lateral hypothalamus (LH) and arcuate hypothalamus (Arc); dashed box denotes area in 20x inset). 3V: $3^{rd}$ ventricle.

(E) Representative image of H129, PRV and double labeling in forebrain (dashed line denotes medial preoptic area (MPA), median preoptic nucleus (MnPO) and anterior commissure area (aca); dashed box denotes area in 20x inset). LV: lateral ventricle; 3V: $3^{rd}$ ventricle.

**Supplemental Table 1. Number of PRV-, H129- and double-labeled neurons in hindbrain, midbrain and forebrain in subdiaphragmatic vagotomized mice with PRV152 injected into the iBAT and H129 injected into the stomach.**

| | PRV152 (# of Neurons) | H129 (# of Neurons) | DOUBLE (# of Neurons) |
|---|---|---|---|
| **Hindbrain** | | | |
| 7N | 2.3 ± 1.1 | 4.7 ± 1.5 | 18.3 ± 7.1*# |
| 12N | 0.7 ± 0.8 | 2.3 ± 1.5* | 10.3 ± 2.3*# |
| Bar | 2.3 ± 0.4 | 7.3 ± 1.1 | 8.7 ± 3.2 |
| Gi | 6.0 ± 1.9 | 20.0 ± 2.8 | 45.0 ± 8.9*# |
| DPGi | 9.0 ± 2.1 | 3.3 ± 1.1 | 20.0 ± 3.1*# |
| GiA | 4.0 ± 1.9 | 2.0 ± 1.4 | 18.3 ± 4.3*# |
| GiV | 12.7 ± 2.9 | 2.7 ± 1.6 | 53.0 ± 11.4*# |
| LPGi | 8.7 ± 2.2 | 5.7 ± 1.5 | 42.3 ± 3.9*# |
| IRt | 13.3 ± 3.9 | 6.3 ± 2.2 | 44.0 ± 10.0*# |
| AP | 1.7 ± 1.1 | 2.0 ± 1.9 | 12.3 ± 4.3*# |
| LC | 18.3 ± 5.8 | 5.0 ± 1.4 | 12.0 ± 2.8 |
| ll | 3.3 ± 1.1 | 9.7 ± 2.3 | 41.3 ± 5.8*# |
| MdD | 5.3 ± 2.3 | 11.3 ± 2.2* | 17.3 ± 1.8*# |
| MdV | 11.0 ± 1.2 | 3.3 ± 2.2 | 43.3 ± 7.1*# |
| Mo5 | 3.3 ± 2.5 | 9.3 ± 3.2 | 11.0 ± 5.0 |
| NTS/Sol | 13.7±3.6 | 9.3±4.1 | 48±10.7*# |
| DMV | 4.7±1.5 | 3.7±0.8 | 5.3±1.5 |
| Parabrachial Areas | | | |
| LPBE | 9.7 ± 2.7 | 0.7 ± 0.4* | 3.0 ± 0.7* |
| LPBS | 7.3 ± 2.9 | 2.3 ± 1.1 | 5.3 ± 1.1 |
| Doral Raphe Areas | | | |
| RMg | 11.3 ± 2.5 | 25.0 ± 5.5 | 128.3 ± 12.1*# |
| ROb | 11.0 ± 4.2 | 0.7 ± 0.8* | 19.0 ± 3.9# |
| RPa | 11.3 ± 1.5 | 6.7 ± 1.1 | 59.3 ± 12.4*# |
| SubCD | 9.7 ± 4.3 | 8.3 ± 2.9 | 38.0 ± 5.0*# |
| SubCV | 4.3 ± 1.8 | 5.3 ± 2.2 | 38.3 ± 7.8*# |
| PPy | 2.7 ± 1.8 | 0.3 ± 0.4 | 37.7 ± 4.3*# |
| rs | 11.0 ± 3.1 | 10.3 ± 2.7 | 32.0 ± 8.9*# |
| KF | 14.7 ± 6.8 | 19.7 ± 5.8 | 51.3 ± 17.4*# |

| | PRV152 (# of Neurons) | H129 (# of Neurons) | DOUBLE (# of Neurons) |
|---|---|---|---|
| **Midbrain** | | | |
| LDTg | 17.0 ± 4.4 | 14.7 ± 0.8 | 37.7 ± 4.3*# |
| ml | 4.7 ± 2.2 | 4.0 ± 1.9 | 22.42 ± 2.25*# |
| pn/PnO | 3.3 ± 1.1 | 2.0 ± 1.2 | 19.3 ± 3.6*# |
| PAG | 74.7 ± 12.5 | 33.3 ± 10.5 | 96.0 ± 31.5# |
| DMPAG | 6.3 ± 1.8 | 1.7 ± 1.1 | 13.7 ± 3.5*# |
| LPAG | 30.3 ± 6.8 | 7.3 ± 1.1* | 36.7 ± 10.5*# |
| VLPAG | 30.19 ± 7.71 | 33.91 ± 9.19 | 21.61 ± 6.23 |
| PPTg | 8.3 ± 1.1 | 4.0 ± 1.9 | 23.0 ± 8.0*# |
| Pa4 | 7.7 ± 1.1 | 0.3 ± 0.4* | 5.7 ± 1.1# |
| DR | 5.0 ± 1.4 | 0.7 ± 0.8* | 4.7 ± 0.8# |
| Su5 | 11.3 ± 3.3 | 11.0 ± 5.6 | 41.0 ± 8.2*# |
| VTA | 5.0 ± 0.7 | 9.7 ± 1.1* | 6.0 ± 1.9 |

| | PRV152 (# of Neurons) | H129 (# of Neurons) | DOUBLE # of Neurons) |
|---|---|---|---|
| **Forebrain** | | | |
| *Hypothalamic* | | | |
| AHA | 1.0 ± 0.7 | 0.7 ± 0.8 | 5.3 ± 0.4*# |
| AHP | 5.7 ± 1.8 | 7.3 ± 1.1 | 9.3 ± 1.1 |
| ADP | 5.0 ± 1.4 | 2.0 ±1.4 | 4.3 ± 2.7 |
| Arc | 15.3 ± 3.3 | 3.7 ± 1.5* | 16.0 ± 2.8# |
| DM | 63.3 ± 9.4 | 30.3 ± 3.3* | 63.0 ± 7.0# |
| LH | 74.3 ± 16.4 | 57.0 ± 10.7 | 132.0 ± 11.1*# |
| PH | 6.0 ± 1.4 | 1.7 ± 1.1* | 17.3 ± 1.8*# |
| LSV | 8.7 ± 2.5 | 0.3 ± 0.4* | 6.3 ± 2.9 |
| Preoptic Areas | | | |
| LPO | 8.0 ± 1.9 | 4.3 ± 1.1 | 26.7 ± 2.9*# |
| MnPO | 6.0 ± 1.9 | 0.7 ± 0.8* | 1.5 ± 0.5* |
| MPA | 23.3 ± 3.2 | 3.7 ± 2.2 | 45.0 ± 15.8# |
| VMPO | 6.3 ± 2.2 | 0.0 ± 0.0* | 2.7 ± 2.2 |
| PVH | 71.0 ± 13.4 | 57.7 ± 2.9 | 181.7 ± 9.6*# |
| PaAP | 11.7 ± 5.7 | 8.7 ± 4.3 | 32.7 ± 13.4 |
| PaLM | 20.7 ± 8.7 | 7.7 ± 4.3 | 41.0 ± 19.9 |
| PaMM | 20.7 ± 9.2 | 18.0 ± 7.8 | 45.0 ± 21.4 |
| PaPo | 14.3 ± 3.6 | 12.0 ± 5.5 | 35.3 ± 12.7# |
| PaV | 3.7 ± 1.1 | 11.3 ± 2.2 | 27.7 ± 8.3*# |
| RCh | 1.7 ± 0.4 | 0.0 ± 0.0 | 2.7 ± 1.1# |
| SCh | 3.3 ± 1.6 | 0.3 ± 0.4 | 6.0 ± 1.4# |
| VMH | 16.0 ± 5.8 | 15.7 ± 1.8 | 28.3 ± 5.0 |
| *Thalamic* | | | |
| SPF | 9.3 ± 3.6 | 1.0 ± 0.7* | 6.3 ± 2.5 |
| PS | 4.7 ± 4.5 | 4.3 ± 3.6 | 9.3 ± 6.6 |
| PSTh | 3.0 ± 1.9 | 1.0 ± 0.7 | 7.7 ± 2.2# |
| ns | 12.3 ± 3.5 | 3.7 ± 2.5 | 21.3 ± 4.3# |

Data are present as means ± SE. *p<0.05 vs. PRV152; #p<0.05 vs. H129

***Hindbrain:*** 7N, facial nucleus; 10N, dorsal motor nucleus of vagus; 12N, hypoglossal nucleus; Bar, Barrington's nucleus; Gi, gigantocellular reticular nucleus; DPGi, dorsal paragigantocellular nucleus; GiA, gigantocellular reticular nucleus, alpha part; GiV, gigantocellular reticular nucleus, ventral part; LPGi, lateral paragigantocellular nucleus; IRt, intermediate reticular nucleus; AP, area postrema; LC, locus coeruleus; ll, lateral lemniscus; MdD, medullary reticular nucleus, dorsal part; MdV, medullary reticular nucleus, ventral part; Mo5, motor trigeminal nucleus; NTS/Sol, nucleus of the solitary tract; DMV: dorsal motor nucleus of the vagus; LPBE, lateral parabrachial nucleus, external part; LPBS, lateral parabrachial nucleus, superior part; RMg, raphe magnus nucleus; ROb, raphe obscurus nucleus; RPa, raphe pallidus nucleus; Sub-CD/V, subcoeruleus nucleus-dorsal/ventral parts; PPy, parapyramidal nucleus; rs, rubrospinal tract; KF, Kolliker-Fuse nucleus.

***Midbrain:*** LDTg, laterodorsal tegmental nucleus; ml, medial lemniscus; PnO, pontine reticular nucleus, oral part; PAG, periaqueductal gray; DMPAG, dorsomedial periaqueductal gray; LPAG, lateral periaqueductal gray; VLPAG, ventrolateral periaqueductal gray; PPTg, pedunculopontine tegmental nucleus; Pa4, paratrochlear nucleus; DR, dorsal raphe nucleus; Su5, supratrigeminal nucleus; VTA, ventral tegmental area.

***Forebrain:*** AHA, anterior hypothalamic area, anterior part; AHP, anterior hypothalamic area, posterior part; ADP, anterodorsal preoptic nucleus; Arc, arcuate nucleus; DM, dorsomedial hypothalamic nucleus; LH, lateral hypothalamic area; PH, posterior hypothalamic area; LSV, lateral septal nucleus, ventral part; LPO, lateral preoptic area;

MnPO, median preoptic nucleus; MPA, medial preoptic area; VMPO, ventromedial preoptic nucleus; PVH, paraventricular nucleus of the hypothalamus; PaAP, paraventricular hypothalamic nucleus, anterior parvicellular part; PaLM, paraventricular hypothalamic nucleus, lateral magnocellular part; PaMM, paraventricular hypothalamic nucleus, medial magnocellular part; PaPo, paraventricular hypothalamic nucleus, posterior part; PaV, paraventricular hypothalamic nucleus, ventral part; RCh, retrochiasmatic area; SCh, suprachiasmatic nucleus; VMH, ventromedial hypothalamic nucleus; SPF, subparafascicular thalamic nucleus; PS,  parastrial nucleus; PSTh, parasubthalamic nucleus; ns, nigrostriatal bundle.
